# The Genetic Basis of Transcriptional Heterogeneity for Basal-Like Features in Pancreatic Ductal Adenocarcinoma

**DOI:** 10.1101/548354

**Authors:** Akimasa Hayashi, Jun Fan, Ruoyao Chen, Yu-jui Ho, Alvin P. Makohon-Moore, Yi Zhong, Jungeui Hong, Hitomi Sakamoto, Marc A. Attiyeh, Zachary A. Kohutek, Lance Zhang, Jinlong Huang, Aida Boumiza, Rajya Kappagantula, Priscilla Baez, Laura D. Wood, Ralph H. Hruban, Lisi Marta, Kalyani Chadalavada, Gouri J. Nanjangud, Olca Basturk, David S. Klimstra, Michael Overholtzer, Christine A. Iacobuzio-Donahue

## Abstract

Pancreatic cancer expression profiles largely reflect a classical or basal-like phenotype. The extent to which these profiles vary within a patient is unknown. We integrated evolutionary analysis and expression profiling in multiregion sampled metastatic pancreatic cancers, finding that squamous features are the histologic correlate of an RNA-seq defined basal-like subtype. In patients with coexisting basal/squamous and classical/glandular morphology, phylogenetic studies revealed that squamous morphology represented a subclonal population. Cancers with squamous features were significantly more likely to have truncal mutations in chromatin modifiers, intercellular heterogeneity for *MYC* amplification, and entosis. These data provide a unifying paradigm for integrating basal-type expression profiles, squamous histology, and somatic mutations in chromatin modifier genes in the context of clonal evolution of pancreatic cancer.

**One Sentence Summary:** Basal type expression profiles in pancreatic cancer have a clonal basis rooted in genetic alterations of chromatin modifier genes.

## Main Text

Despite the wealth of data pertaining to the biology and genetics of pancreatic ductal adenocarcinoma (PDAC), this solid tumor remains one of the most lethal tumor types (*1–3*). Large scale sequencing studies have revealed the recurrent genomic features of this disease that target a defined number of core pathways (*4–8*). In some patients a genome instability signature is also seen based on either microsatellite instability or on a high number of structural rearrangements (*5, 9*). Transcriptional studies have revealed that PDAC can be segregated into two major subtypes termed “classical” and “basal-like” (*6, 7, 10, 11*).

We previously leveraged multi-regional sampling to define the genetic evolution of pancreatic cancer metastasis. We found within each patient that the primary tumor and metastases shared identical driver gene mutations, suggesting that at least one major clonal sweep had occurred. The cells that descended from this sweep were endowed with all of the genetic drivers needed to metastasize (*6, 7, 11*). We have also observed that metastases from these same patients may have divergent morphologic and molecular features despite identical genomes (*12*). In light of these observations we posited that an integrated analysis of the histologic, genomic and transcriptional features of PDAC would provide insight into tumor progression, both within the primary and metastatic sites.

## Results

We reviewed hematoxylin and eosin stained sections prepared from more than 7,000 unique formalin-fixed paraffin-embedded tissues from 156 research autopsy participants spanning two institutions, all of whom had been clinically or pathologically diagnosed with PDAC premortem. After histologic review 33 cases were excluded (see Fig 1.A., Fig. S1A and Data S1.) leaving 2944 individual sections from 123 cases (median 17 tumor sections per case) that fulfilled our criteria for further study. Histologic review in combination with immunohistochemical labeling of representative blocks for the common squamous differentiation markers CK5/6 and p63 (*13, 14*) was performed so that each individual formalin-fixed section was categorized as having a conventional glandular pattern of growth (GL), squamoid features (SF), or squamous differentiation (SD) (Fig. 1B, Fig. S2). Of 2944 blocks, 490 (16.6%) showed squamoid features (SF) or squamous differentiation (SD) (Fig 1C). As described in previous studies (*14*) SF/SD occurred as discrete regions within a PDAC or as an admixture of glandular and squamous morphologies. We therefore estimated the proportion of squamous differentiation in each carcinoma based on the number of blocks with SF/SD and the area of squamous differentiation within each block. Seven PDACs (5.7%) met WHO criteria (*15*) for adenosquamous carcinoma (ASC), six PDACs (4.9%) had focal (<30%) squamous differentiation and two PDACs (1.6%) had squamoid features (Fig. 1D and Fig. S1B). Five PDACs (PAM02, PAM22, PAM28, PAM55, PAM80) had all three morphologies present (Fig. 1D and Fig.1E). When present, the proportion of SF/SD in a carcinoma ranged from 2% to 80% (Fig.1D). By univariate analysis patients with ASCs or PDACs with SF/SD had a poorer survival than did patients with PDACs without SF/SD (Fig. 1F), compatible with previous findings (*16*). The prevalence of ASC in this cohort of 129 patients with end stage disease is higher than that reported for surgically resected tumors (5.7% of our cohort vs 0.9% in Boyd et al. P < 0.0001, Chi-square test) (*16*) indicating that SF/SD develops in association with tumor progression.

**Fig. 1.**
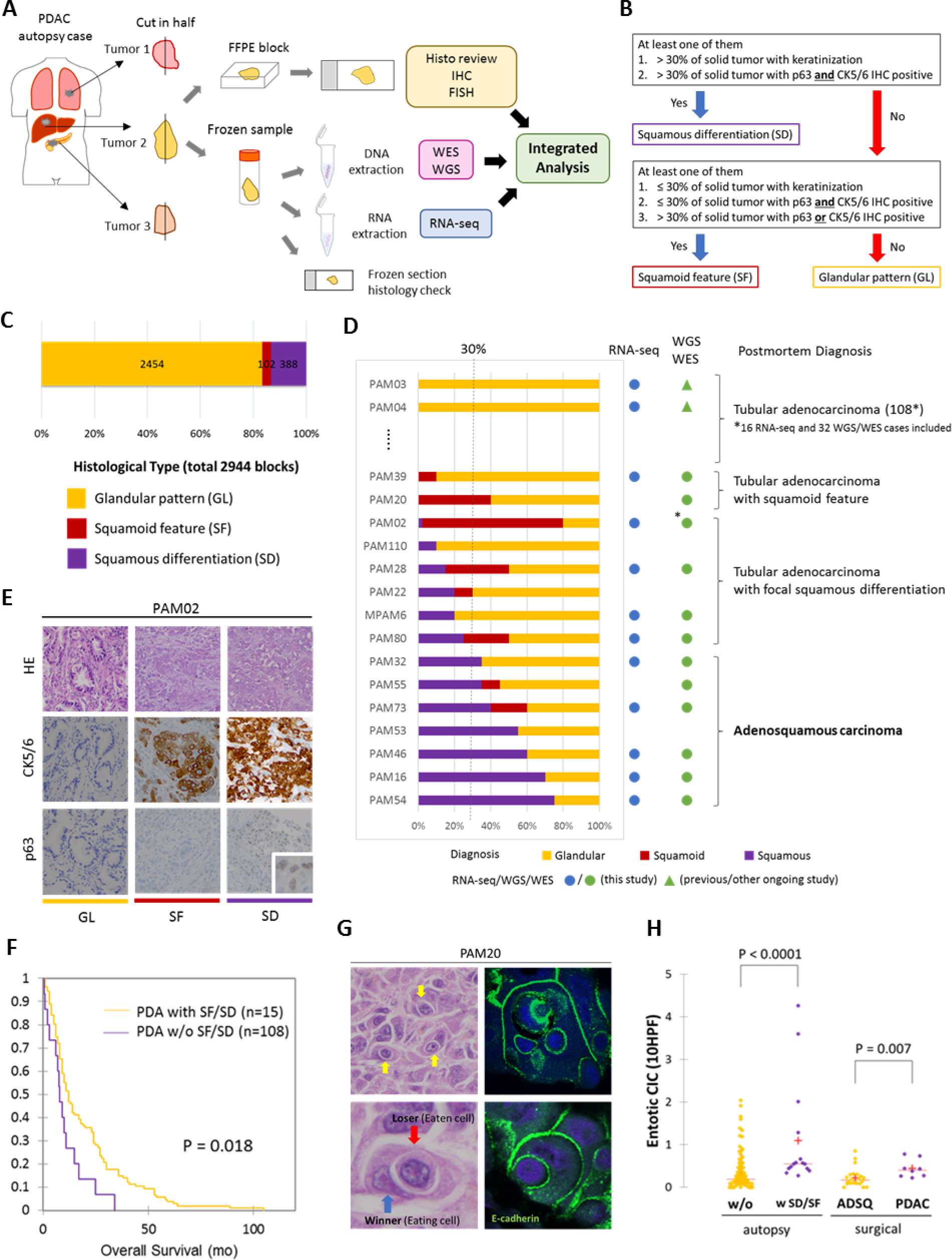
Study Overview and Morphologic Heterogeneity for Squamous Features in Pancreatic Ductal Adenocarcinoma. (A) Study overview of integrated analysis in pancreatic ductal adenocarcinoma using multiregional sampling. **(B)** Schematic for classification of sections. Squamous differentiation or squamoid feature was determined for each block in all cases based on the combination of histomorphologic features and p63 and CK5/6 immunohistochemistry (IHC). **(C)** Summary of block diagnoses. **(D)** Postmortem case diagnoses based on combination of the number of blocks with SF or SD of all blocks analyzed per patient and the percent of SF or SD within each positive block. *PAM02 was reanalyzed for this study using previous data. **(E)** Representative histomorphologic and immunocytochemical images of glandular pattern (GL), squamoid feature (SF) and squamous differentiation (SD) in patient PAM02. SD areas showed solid growth pattern with both CK5/6 and p63 positivity, while SF areas showed CK5/6 positive labeling but are negative for p63. **(F)** Kaplan-Meier analysis of PDAC with or without SF/SD. PDAC with SF/SD showed poorer prognosis than PDACs without SF/SD. **(G)** Representative histomorphologic and immunofluorescent images of entotic CIC in patient PAM20. A clearly defined ‘moonshape’ host nucleus, intervening vacuolar space and internalized cell is identified. Immunofluorescent images clear e-cadherin membranous labeling of the winners (eating cells) and losers (eaten cells). **(H)** Average number of entotic cell in cell structures (entotic CIC) in PDAC with or without SF/SD.

Based on histologic review of all 2944 sections we also noted that ASCs and PDACs with SF/SD exhibited entosis, a distinct form of cell death in which one cancer cell engulfs another (Fig. 1G) (*17*). To more rigorously determine the relationship of SF/SD to entosis we adopted strict criteria to count entotic cell-in-cell structures (CIC) (see Methods) (*18*). The number of entotic CIC was higher in PDACs with SF/SD or ASCs compared to PDACs without SF/SD in our cohort (1.095±1.240 versus 0.365±0.437 per 10HPF respectively, P < 0.0001, Mann–Whitney U test). To determine if entosis is more reflective of stage of disease versus morphology, we reviewed an independent cohort of 30 resected PDACs that included eight ASCs. Similar to the findings in the autopsy cohort, resected ASCs had more entotic CIC than conventional PDACs (0.441±0.213 versus 0.226±0.219, respectively, P = 0.007, Mann–Whitney U test) (Fig. 1H). However, there was no difference in the number of entotic CICs in autopsy PDACs with SF/SD or ASCs compared to surgically resected ASCs, suggesting entosis is a feature of SF/SD specifically.

We next sought to determine the extent that the observed morphologic findings correspond to the “classical” and “basal-like” type transcriptional signatures described (*7, 11*). We extracted total RNA from 480 frozen samples in triplicate; in all cases the frozen tissue was matched to the formalin-fixed sections used for morphologic and immunohistochemical analyses. A total of 214 frozen samples from 27 patients in our cohort (median 6 samples, range 1 to 26 samples per patient) meeting quality criteria (see Methods) were used for RNA sequencing (Data S2). These 27 cases included five ASCs and five PDAC with focal SF/SD; for these 10 cases the GL and SF/SD regions were independently extracted and analyzed. Normalized mRNA expression levels of *TP63*, *KRT5* and *KRT6A* confirmed that GL-PDAC samples had the lowest expression of all three markers whereas SD-PDAC samples had the highest expression levels of all three markers. SF-PDACs had an intermediate expression pattern between GL-PDACs and SD-PDACs (Fig. S3). Consistent with this finding, network analysis highlights *KRT5* and *KRT6A* as “hub” genes in samples with SF/SD morphology, and SF/SD morphology shows more complex co-expression patterns in keratin filament & keratinization pathways than in samples with GL morphology. We next classified our 214 samples into “classical” and “basal-like” PDAC subtypes using the 50 pancreas cancer gene set reported by Moffitt et al. (*11*) because recent TCGA re-analysis showed that this classification was least affected by tumor purity or stromal contamination (*7*). This revealed an almost perfect concordance of morphologic features with transcriptional subtype, as most SF-PDAC and all SD-PDAC samples corresponded to the “basal-like” expression pattern, whereas most GL-PDACs corresponded to the “classical” type pattern (Fig. 2A). Principal component analysis using this same gene set revealed a similar distribution based on morphologic features or RNA expression subtype, whereas no relationship was found for site of harvesting of each sample (primary or metastasis) (Fig. 2B). This confirms that the “basal-type” expression signature as defined by this 50 gene signature reflects squamous differentiation in PDAC.

**Fig. 2.**
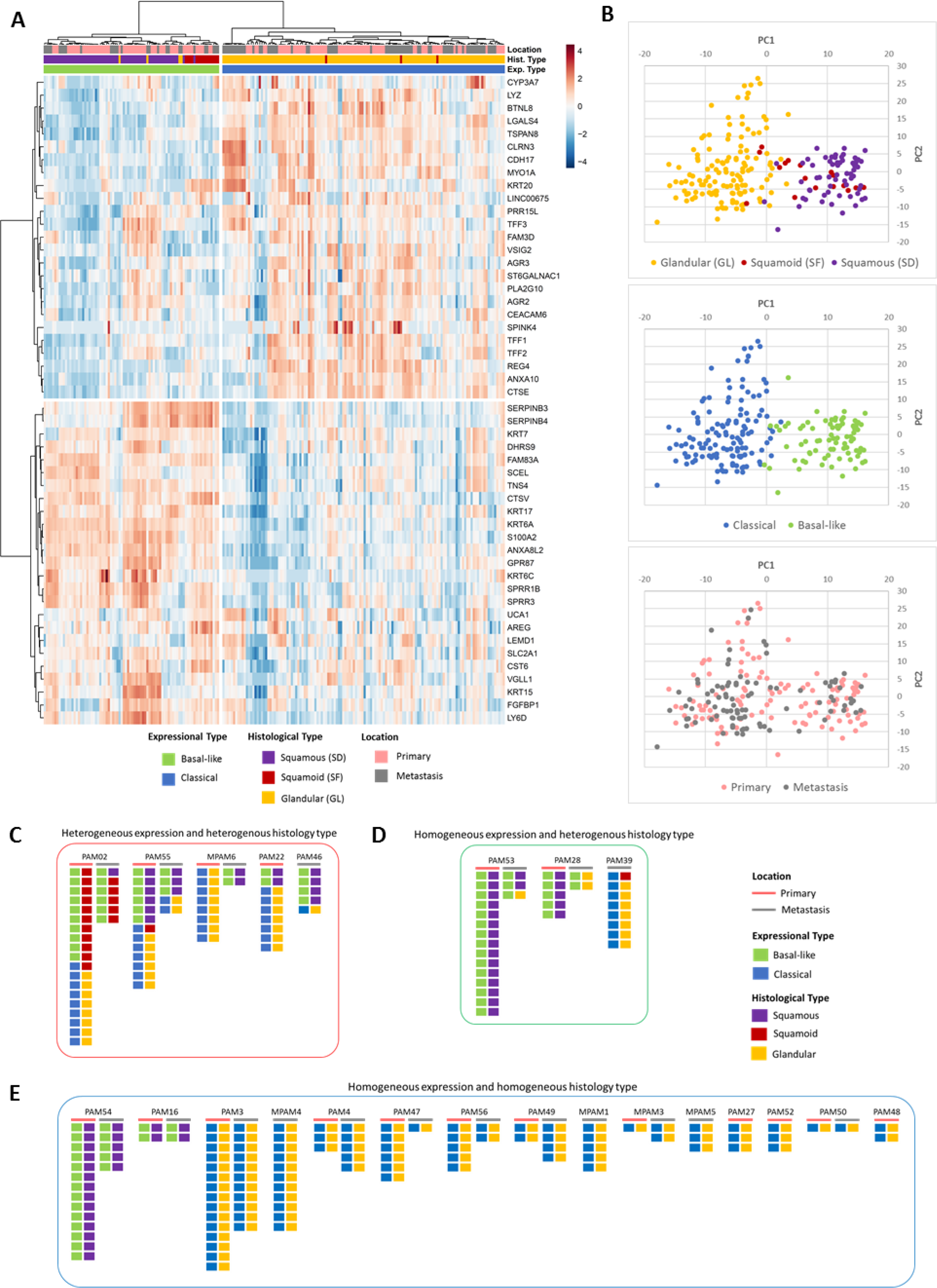
Transcriptional Heterogeneity for Squamous Features in Pancreatic Ductal Adenocarcinoma. RNA sequencing (RNA-seq) of was performed on snap-frozen tissues of 214 unique samples from 27 patients including five with ASC and five with focal SF/SD. RNA-seq data were used to classify each of the 214 samples into “basal-like” and “classical” tumors (Moffitt et al. Nature Genetics, 2015). Both the heatmap **(A)** and PCA plots **(B)** indicate a strong correlation of SF/SD morphology with a “basal-like” transcriptional signature, and GL morphology with a “classical” transcriptional signature. **(C)-(E)** Integrated analysis of transcriptional subtype with unique block diagnosis indicates intratumoral heterogeneity for both transcriptional signatures and histomorphologic features in a subset of PDACs.

For 23 of these 27 patients two or more samples were analyzed by RNA-seq. Intratumoral heterogeneity for expression profiles was identified in five patients (PAM02, PAM22, PAM46, PAM55, MPAM6) indicating that the “classical” and “basal-like” subtypes can co-exist within a single patient (Fig.2C). With two exceptions (one primary tumor sample each in PAM02 and PAM55) the transcriptional signatures correlated with the histologic features of the sample. In a separate set of three patients (PAM28, PAM39, PAM53) all samples analyzed were homogenous for their transcriptional subtype despite a degree of morphologic heterogeneity (Fig. 2D). These included a “basal-like” transcriptional signature but glandular morphology in the metastases of PAM28 and PAM53, and a “classical” expression signature in a metastasis with squamoid features in PAM39. Finally, in 15 patients all samples studied were homogeneous with respect to both their transcriptional subtype and morphologic pattern (Fig. 2E). The majority of these cases had a glandular morphology (PDAC-GL) and a “classical” type expression signature, although in two patients (PAM16, PAM54) prominent squamous differentiation was identified in all samples analyzed for the tumor and had a “basal-like” expression signature.

We next determined the relationship of the coding genomic landscape to the presence of SF/SD by performing multiregion whole exome or whole genome sequencing on frozen samples matched to histologically and immunohistochemically characterized formalin-fixed sections in 43 patients. Overall the genetic features of this cohort were consistent with the PDAC genomic landscape (Fig. 3, Data S4)(*4–8*) and no mutations of *UPF1* were identified that have previously been reported in ASC (*19*). However, two carcinomas had a *KDM6A* mutation (*6, 20*), both in females and one with an ASC, leading us to more closely evaluate all chromatin modifier gene mutations identified in these 43 patients. The most common chromatin modifier gene with a deleterious mutation was *ARID1A* (four carcinomas, 9%), followed *KMT2D and KMT2C* (three carcinomas, 7%), *ARID2*, *KDM6A, SMARCA4* (two carcinomas each, 5%). With one exceptional case the mutations in chromatin modifier genes were mutually exclusive (Fig. 3). Six of 12 patients (50%) with a PDAC with SF/SD morphology or ASC had a mutation in a chromatin modifier gene compared to 9 of 31 patients (29%) with a PDAC with GL morphology, a difference that did not reach statistical significance (P = 0.287, two-sided Fisher Exact Test). We also noted *RB1* mutations in three PDAC with SF/SD cases (25%) compared to only one case without SF/SD (3%), although this finding was also not statistically significant (P = 0.059, two-sided Fisher Exact Test). To better understand this finding we compared the frequency of somatic alterations in chromatin modifier genes in our dataset to that in the MSK Clinical IMPACT cohort that is highly enriched for treatment naïve PDAC (*21*). This revealed a statistically significant enrichment of chromatin modifier mutations in our cohort of patients with end stage disease (P = 0.017, two-sided Fisher Exact Test). *RB1* alterations were also more frequent in our end stage PDAC cohort compared to MSK-IMPACT (P = 0.017, two-sided Fisher Exact Test). *RB1* has recently been shown to play a role in telomeric chromatin architecture (*22*).

**Fig. 3.**
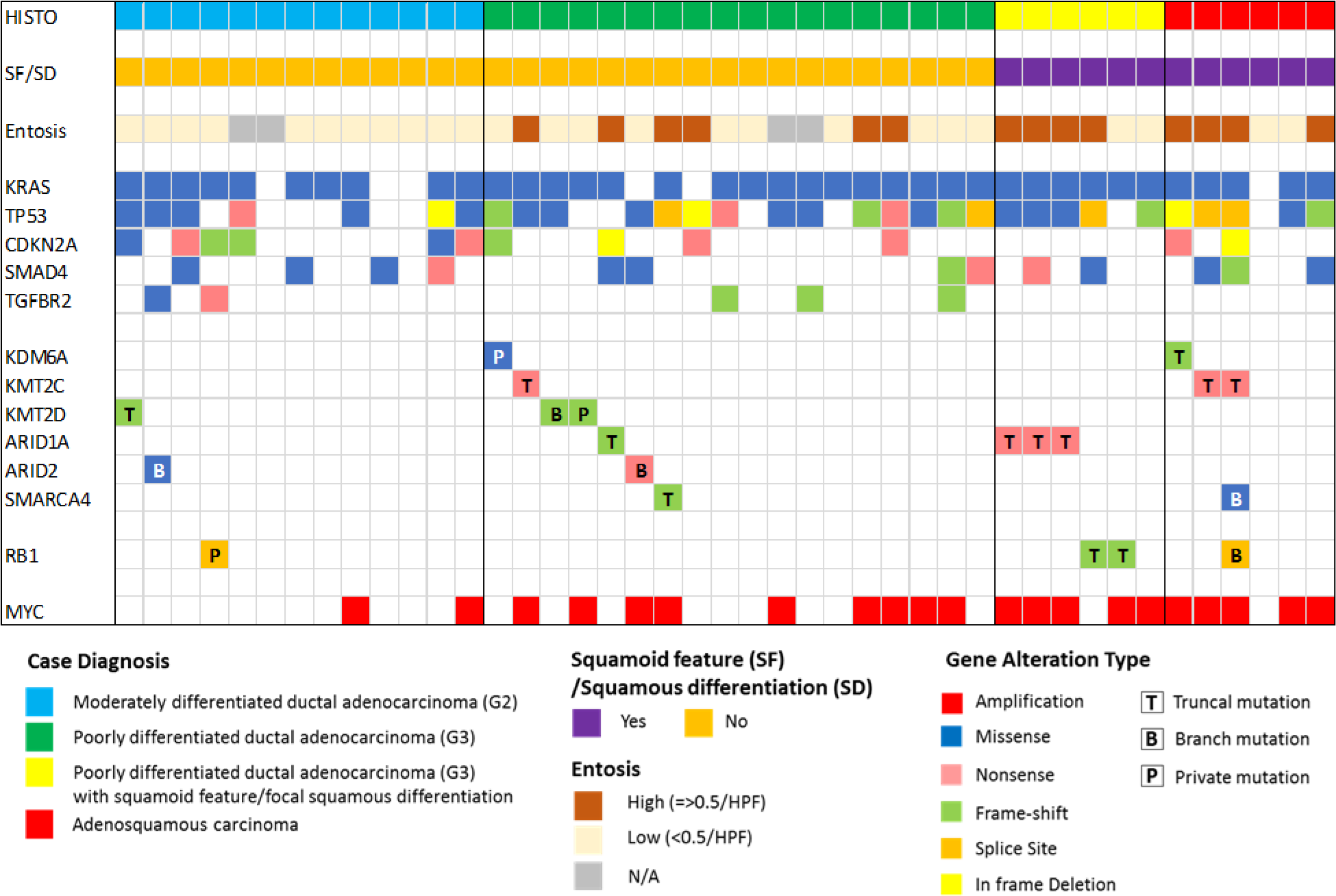
Genomic Landscape of End Stage Pancreatic Ductal Adenocarcinomas with and without Squamous Features. Oncoprint illustrating the driver gene somatic alterations of 43 cases with respect to their histologic and immunolabeling profiles. Truncal mutations in chromatin modifier genes and MYC amplification are significantly enriched in PDACs with focal SF/SD and ASCs.

High quality single nucleotide variants and small insertions/deletions identified for each sample were used to recreate the phylogenetic relationships among the spatially distinct samples within each patient. While there was no difference in the prevalence of mutations in chromatin modifier genes in cancers with or without SF/SD, these analyses indicated an influential relationship among the evolutionary timing that a mutation in a chromatin modifier gene arose and the extent of squamous morphology in the carcinoma. For example, six of six chromatin modifier gene mutations identified in PDACs with SF/SD morphology or ASCs were truncal in origin (Fig. 3, Fig. 4, Fig. S3-S8), compared to only four of the ten mutations in PDACs with GL morphology (P = 0.034, two-sided Fisher Exact Test). In the remaining PDACs with GL morphology, mutations in chromatin modifier genes were assigned to a branch or were private to a single sample in that patient. Curiously, we noted two PDACs with SF/SD and wild type chromatin modifier genes (PAM28, MPAM6) had deleterious truncal mutations of *RB1* (Fig. S9, Fig. S10). Collectively, we conclude that transcriptional heterogeneity for “basal-like” features corresponds to morphologic heterogeneity for SF/SD, and these features occur in the setting of truncal mutations in chromatin modifier genes, most often but not exclusively *ARID1A, KMT2C or KMT2D*.

**Fig. 4.**
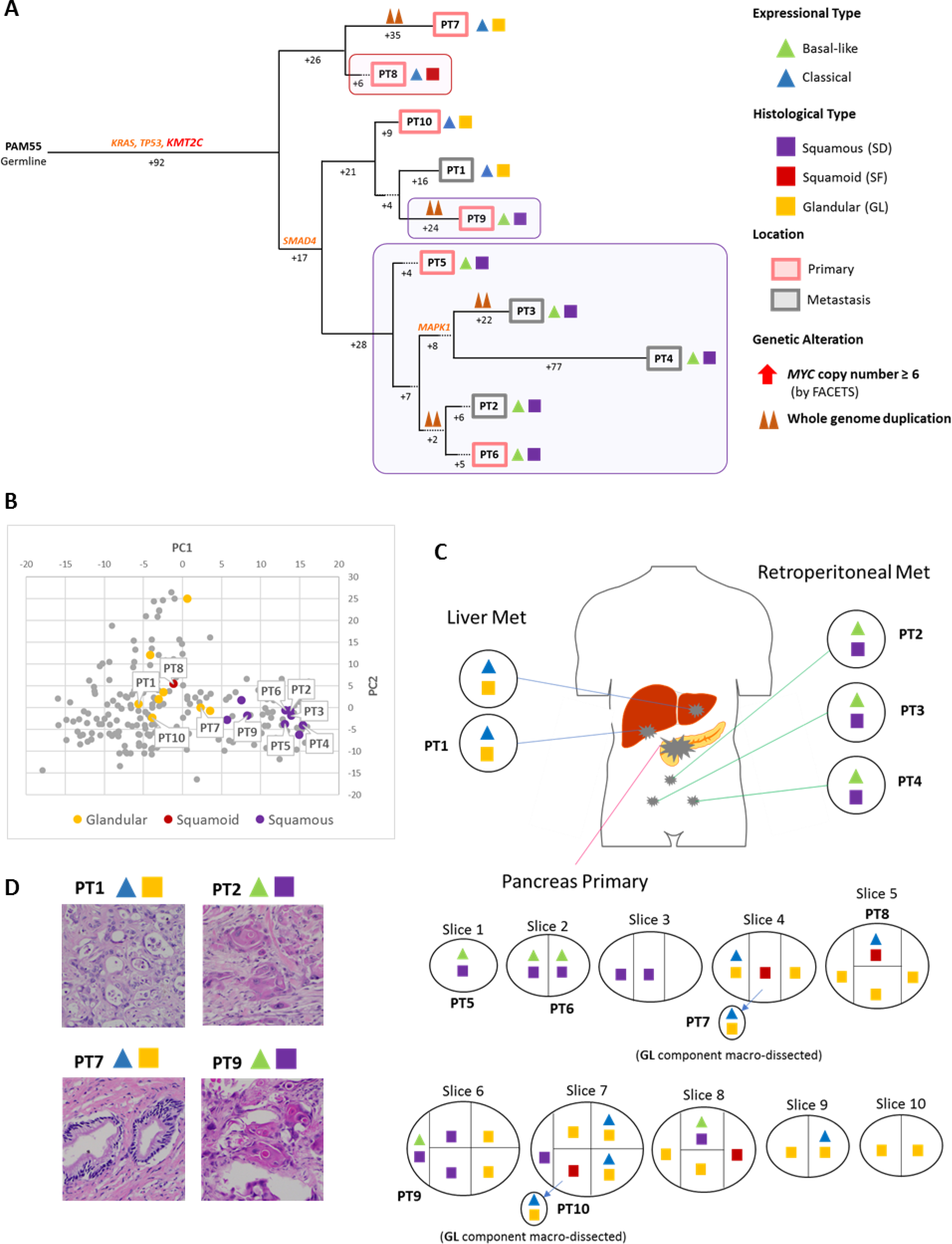
Integration of Transcriptomic and Morphologic Features with Phylogenetic Patterns in Pancreatic Ductal Adenocarcinoma PAM55 with a Truncal Chromatin Modifier Mutation. **(A)** Phylogenetic tree of patient PAM55. Truncal driver genes are notable for a *KMT2C* somatic alteration. Red and purple outlines indicate samples that have SF/SD based on RNAseq (triangles) and/or histology (squares). The predicted timing of somatic alterations in driver genes and whole genome duplication are also shown. Mutations in chromatin modifier genes are in red font, all others in orange. SF/SD in this carcinoma have arisen as three independent subclones as defined by their genetic features: in primary tumor sample PT8, in primary tumor sample PT9, and in the subclone giving rise to the evolutionary related primary tumor samples PT5 and PT6 and metastases PT2-PT4. **(B)** Principal Components Analysis illustrating intratumoral expressional heterogeneity and the transition between GL and SF/SD. **(C)** Relationship of anatomic location to morphologic and transcriptional heterogeneity. Shown is the spatial location of each sample within the primary tumor or distant sites and their corresponding transcriptional and histological subtypes. Both histologic and transcriptional heterogeneity are identified in the primary tumor whereas retroperitoneal metastases showed SD with “basal like” type expression and multiple liver metatases showed GL with “classical” type expression. **(D)** Representative histologic images of tumors in this same patient.

We next determined the relationship of heterogeneous morphologic or transcriptional features to the derived phylogenetic relationships of spatially distinct samples within a single patient. In 10 of 12 patients the squamous feature (SF/SD) component was a clonal population, i.e. all samples with SF/SD were phylogenetically more closely related to each other than to the sample(s) with GL morphology in the same patient (Fig. 4, 5 and Fig. S6, S8-S10, S12, S13). These phylogenetic relationships did not imply a shared anatomic location, as genetic, morphologically and transcriptionally similar samples could be found in both the primary tumor and in metastatic sites. In the remaining two patients (PAM22 and PAM39) the SF/SD was exclusive to a single sample analyzed (Fig.S7, Fig. S11). The integration of phylogenetic trees, morphologic features and spatial location also suggested that SF/SD can develop independently in the same neoplasm, for example in PAM55 (Fig. 4) in which samples PT8, PT9 and samples PT2-PT6 were contained within three different clades respectively. This suggests that beyond truncal genetic alterations in chromatin modifier genes, subclonal populations with SF/SD may be further defined by a combination of epigenetic and/or microenvironmental cues (*12, 23*).

**Fig. 5.**
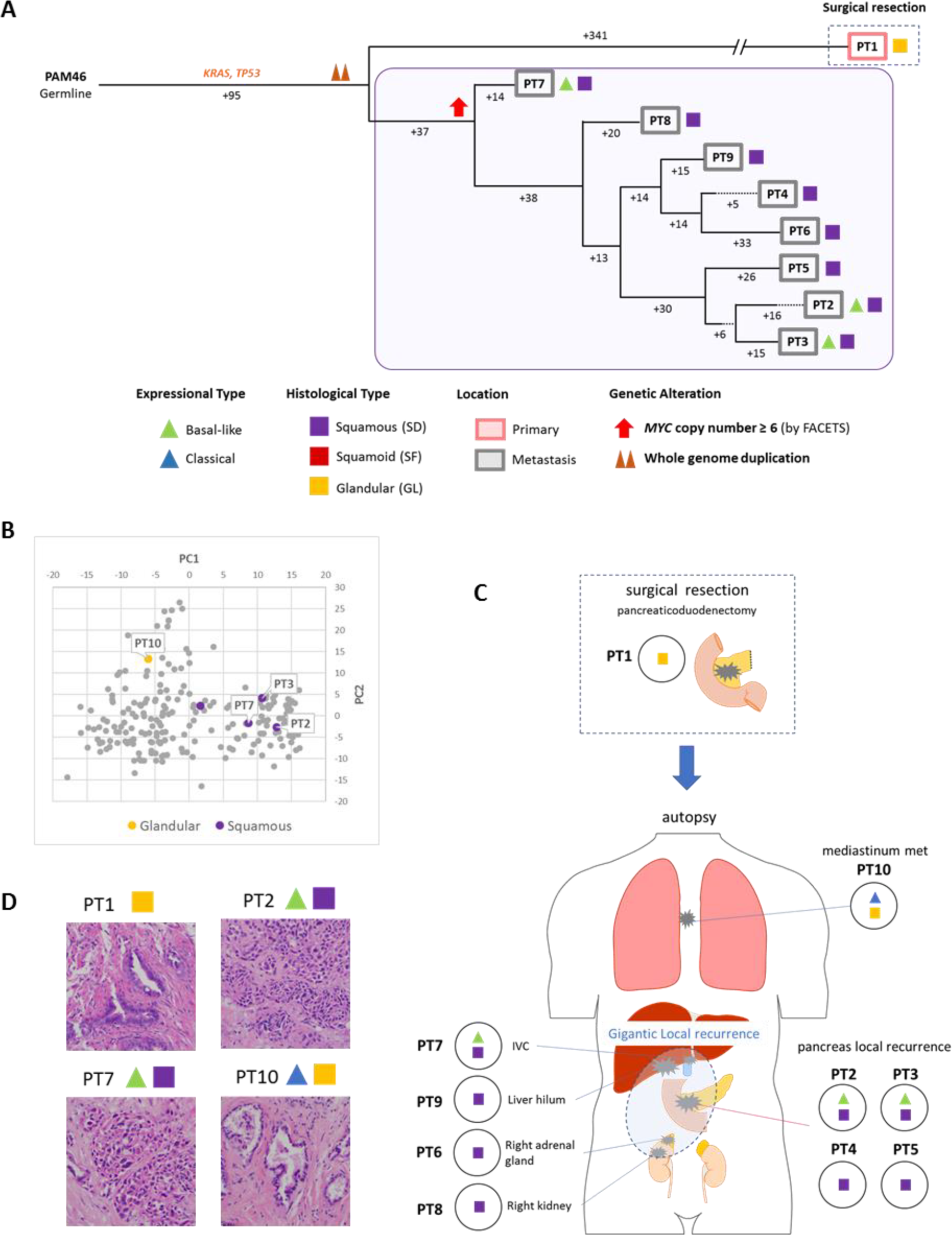
Integration of Transcriptomic and Morphologic Features with Phylogenetic Patterns in Pancreatic Ductal Adenocarcinoma PAM46 with *MYC* amplification. **(A)** Phylogenetic tree of patient PAM46. Purple outline indicates samples that have SF/SD based on RNAseq (triangles) and/or histology (squares). The predicted timing of somatic alterations in driver genes, whole genome duplication and *MYC* amplification are also shown. No mutations in chromatin modifier genes were identified. *MYC* amplification (>6 copies) was detected in all samples of the local recurrence but not the original resected primary tumor PT1. **(B)** Principal Components Analysis indicates PT10 shows a different expressional type from other samples. **(C)** Relationship of Anatomical Location to Morphologic and Transcriptional Heterogeneity. Shown is the spatial location of each sample within the primary tumor or distant sites and their corresponding transcriptional and histological subtypes. GL pattern was only identified in primary surgical resection and mediastinum metastasis (PT10) at autopsy.**(D)** Representative histologic images of representative tumors in the same patient.

To gather insight into potential molecular features that contribute to the development of SF/SD in PDAC, we mined our RNAseq dataset to determine the transcriptional differences between samples with GL morphology and SF/SD morphology in an unbiased manner. Gene set enrichment analysis (GSEA) using Hallmark genesets and transcription factor target genesets (Methods, Data S5) revealed MYC target gene expression as significantly enriched in samples with SF/SD compared to GL morphology (Fig. 6A, Data S6, S7), a finding similar to that reported by Bailey et al. (*6*). To further determine the significance of this observation we reviewed our RNAseq data specifically for *MYC* gene expression. This revealed a significantly higher *MYC* transcript abundance in samples with SF/SD morphology compared to those with GL morphology (Fig. 6B). As *MYC*, located in chromosome 8q 24.21, is a known target of amplification in PDAC (*24, 25*) we performed fluorescent in situ hybridization (FISH) analysis for *MYC* copy number in eight carcinomas wherein both GL and SF/SD morphologies were present within the same tumor/section. In all eight examples *MYC* copy number was significantly higher in regions with SF/SD morphology compared to regions with GL morphology (Fig. 6C-D). We next more broadly analyzed *MYC* amplification across our cohort. *MYC* amplification was significantly associated with squamous subtype (amplification observed in two of 13 Grade 2 PDACs, nine of 18 Grade 3 PDACs, 10 of 12 PDACs with SF/SD or ASCs, P = 0.003, two-sided Fisher Exact Test). *MYC* amplification was also more prevalent in this cohort compared to the reported prevalence in resectable disease (21/43 vs 5/149 in TCGA, P < 0.0001, two-sided Fisher Exact Test) (*5, 7*) as well as significantly correlated with poor outcome (Fig. 6E) and a high number of entotic CICs (P = 0.019, two-sided Fisher Exact Test) (Fig. 3). Overall these findings indicate that gains of *MYC* copy number are correlated with PDAC progression, and particularly so with SF/SD.

**Fig. 6.**
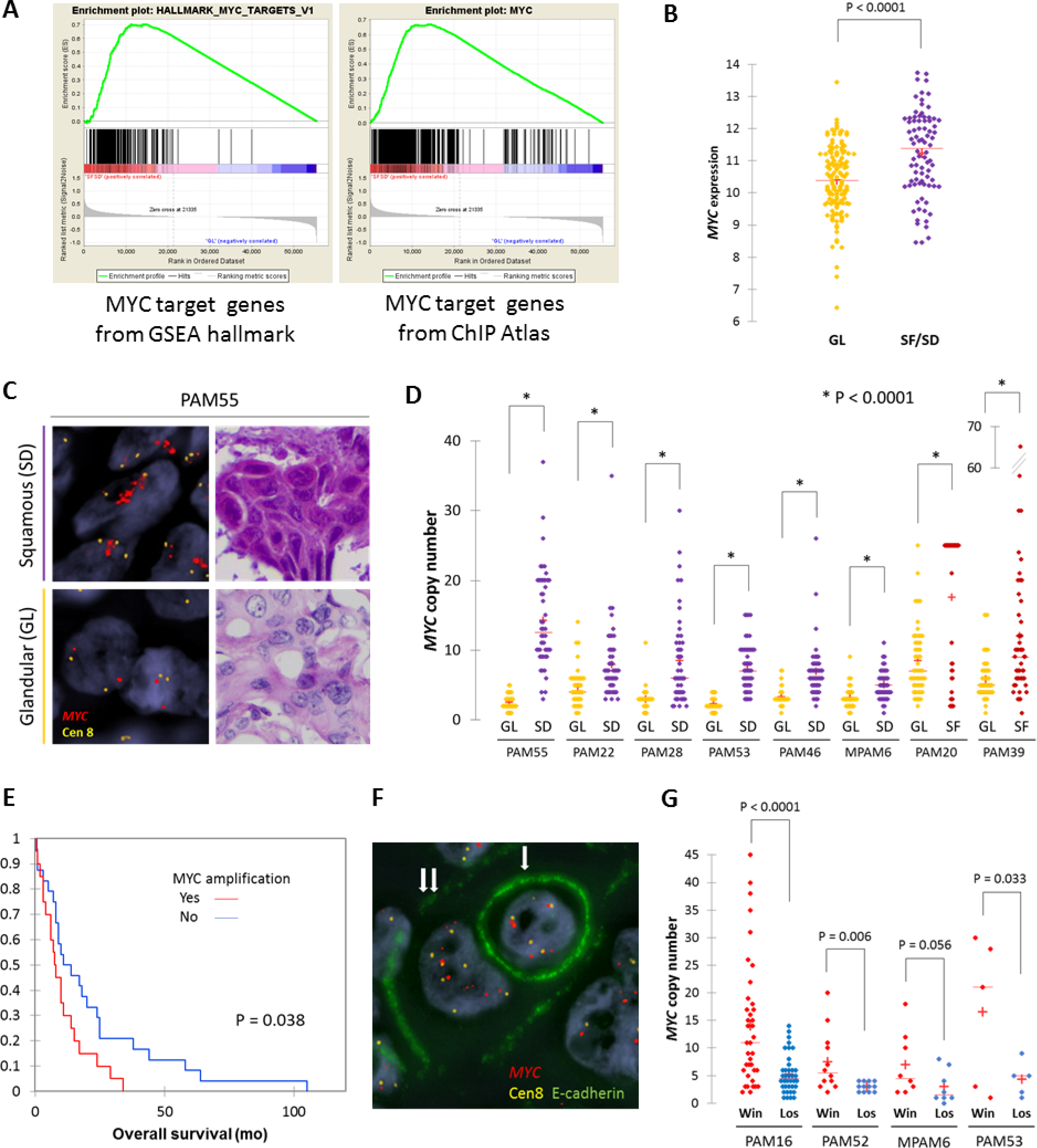
Squamous Features in Pancreatic Ductal Adenocarcinoma Correspond to Enhancement of *MYC*. **(A)** Gene-set enrichment analysis (GSEA) using hallmark gene sets and transcription factor target gene sets collected from ChIP-Atlas identify MYC target genes as the top ranked gene set in SF/SD (see also Supplemental Tables 6 and 7). **(B)** Normalized *MYC* RNA transcript abundance is significantly higher in SF/SD samples than in GL samples. **(C)** Representative images of *MYC*-FISH analysis in SF/SD and GL regions. **(D)** Analysis of *MYC* copy number in eight cases indicates that *MYC* is significantly amplified in SF/SD regions compared to GL regions in the same carcinoma. (**E)** Kaplan-Meier analysis indicating patients whose carcinomas have *MYC* high (>= 6) copy number have a worse outcome than carcinomas with low *MYC* copy number. (**F)** Representative images of entosis (single arrow: loser (eaten cell), double arrows: winner (eating cell)). **(G)** Winner cells have higher *MYC* copy number than loser cells.

In light of the correlation of both *MYC* amplification and entosis with SF/SD we more closely determined the relationship, if any, between these two observations by reviewing nine cases with concurrent *MYC* amplification and entotic CICs. Specifically, we determined *MYC* copy number in matched winner cells (eating) and loser cells (eaten) (Fig. 6F). This revealed a remarkable degree of intercellular heterogeneity for *MYC* copy number in that winner cells had 8.3 ± 10.5 copies of *MYC* compared to only 2.4 ± 3.2 copies per loser cell (P < 0.0001, Mann-Whitney U-test) (Fig. 6G). After normalization for chromosome 8 copy the winner cells retained a higher copy number compared to loser cells (2.1 ± 1.4 copies per winner cell compared to 1.7 ± 0.9 copies per loser cell), but the difference was not statistically significant (P = 0.103, Mann-Whitney U-test) suggesting that gain of *MYC* copy number is selected for in the context of gains in ploidy (*26*). We therefore evaluated the approximate timing of *MYC* copy number gain during clonal evolution based on FACETS copy number and ploidy estimations generated for the 12 sequenced cases for which phylogenies were derived*. MYC* amplification was present in five cases, all in a subclonal manner (Fig. 5, Fig. S6-S8, S11). All five cases had whole genome duplication in one or more samples, and in three cases the phylogenies indicated that *MYC* amplification accompanied or followed gains in ploidy (Fig. 5, Fig. S7, S11). Our integrated phylogenetic analyses and morphologic studies further indicated that in four cases the samples with SF/SD occurred in a lineage derived from the subclonal population with *MYC* amplification (Fig 5, Fig S6-S8). Together, these findings buttress the notion that *MYC* amplification contributes to the development of SF/SD in PDAC.

## Discussion

We describe a unifying paradigm for transcriptional subtypes, squamous morphology and somatic mutations in chromatin modifier genes that is rooted in phylogenetic analyses. These insights provide the context in which to understand the significance of these molecular events for more rigorous stratification of PDAC patients for personalized medicine approaches.

While mutations in *ARID1A, KMT2C* and related genes have consistently been identified in large scale screens of the PDAC genome (*6, 7*), their significance for the natural history of PDAC has remained unclear. We now show that the evolutionary context in which these mutations occur is related to the likelihood the PDAC will develop squamous morphology. This likelihood is not absolute, as evidenced by the deceased patients in our cohort with poorly differentiated PDACs with truncal mutations in chromatin modifier genes. While our findings are consistent with reports that ASCs are associated with a worse outcome (*16*), they contradict those that report an improved outcome in PDACs with mutations in *ARID1A, KMT2C* and related chromatin modifier genes (*27, 28*). Future efforts that consider somatic mutations in these genes specifically in the context of whole genome duplication, *MYC* copy number and morphologic features may resolve this discrepancy.

Our data also illustrate that glandular and SF/SD morphologies, and by extension the classical-type and basal-type expression signatures, coexist in the same PDAC. While prior studies of ASC have also reported this phenomenon (*14, 29, 30*), we now show that the SF/SD component arises from a subclonal population. This raises two possibilities for understanding SF/SD. First, SF/SD may develop from classical-type gland forming pancreatic cancer. The paucity of data reporting small, early stage ASCs and that SF/SD are commonly found in association with conventional glandular features are consistent with this possibility (*29*). Moreover, whereas we found that SF/SD may arise during the clonal evolution of a PDAC we did not observe the converse scenario by phylogenetic analysis, i.e. a subclonal glandular component arising in a predominant SF/SD neoplasm. While we believe the former is the most parsimonious explanation, we acknowledge a second possibility where a common phenotypic intermediate cell type gives rise to both classical-type and basal-type phenotypes. Our study relied on bulk and macrodissected tissues thus we did not reach the level of resolution required to answer this question definitively.

These data also contextualize the significance of *MYC* copy number gains in PDAC by illustrating it is selected for during tumor progression and in association whole genome duplication. Furthermore, we identify an unappreciated feature of *MYC* in PDAC, intercellular heterogeneity for copy number that is associated with entosis. Entosis, a process in which a cancer cell engulfs its neighbor, represents a form of cell competition that stimulated by low glucose environments(*17, 31*). Intriguingly, *MYC* expression has also been shown to promote competition between normal cells in both fly and mammalian tissues during development (*32, 33*) suggesting a new potential mechanistic parallel between intercellular heterogeneity for *MYC* copy number and stimulation of cell competition. In PDAC specifically, these observations provide clues to the microenvironmental changes, i.e. glucose depletion, that contribute to the development of SF/SD in association with mutations in chromatin modifier genes (*34*).

We expect that our findings will have implications for understanding other solid tumor types as well in which these mutations occur and/or that develop squamous features in the course of disease progression. Ultimately, our hope is that comprehensive studies such as this pave the way for identifying novel therapeutic vulnerabilities or re-evaluation of the utility of currently available therapies based on the genotypes and phenotypes assessed.

## Acknowledgments

We are grateful to Gokce Askan and Jacklynn V. Egger for assistance in identifying resected adenosquamous samples for use in this study, and to Shinya Oki for technical support. We gratefully acknowledge the members of the Molecular Diagnostics Service in the Department of Pathology for MSK IMPACT.

## Author contributions

A.H., J.F. and C.I.-D. designed the study; A.H., J.F., A.P.M.-M, H.S., M.A.A., A.B., R.K., P.B., L.D.W., R.H.H., C.I.-D. collected autopsy samples; A.H. and C.I.-D. reviewed histology of autopsy samples and selected cases; O.B., D.K., A.H. and C.I.-D. reviewed pathology of surgical cases; A.H., R.C., M.O., G.J.N. and C.I.-D. reviewed entosis of Immuno-FISH slides; A.H. and J.F. prepared RNA samples; A.P.M.-M., J.H., H.S., Z.K. and A.H., prepared the DNA samples; A.H., Y.Z. and C.I.-D. performed RNA sequencing; Y.H., A.H., L.Z. and J.H. analyzed RNA sequencing results; A.P.M.-M., J.H., Z.K., H.S. M.A.A., A.H., and C.I.-D. performed DNA sequencing; A.P.M.-M., M.A.A., J.H., A.H. and C.I.-D. analyzed DNA sequencing results and derived the phylogenies; L.M., K.C. and G.J.N. performed Immuno-FISH; R.C. performed knockdown experiments; A.H., R.C., Y.H., M.O. and C.I.-D. wrote the manuscript; all authors reviewed and edited the final manuscript.

## Data and materials availability

Sequence data have been deposited at the European Genomephenome Archive (EGA) under accession number EGAS00001002580.

## Supplementary Materials

### Other Supplementary Materials for this manuscript include the following

Data S1. Clinical information of PDA with squamous differentiation or squamoid feature.

Data S2. Sample information of RNA-seq, WES, WGS and target-seq.

Data S3. RNA expression, released after publication

Data S4. Gene alternation of our cohort and MSK IMPACT Clinical Sequencing Cohort.

Data S5. Transcription factor (TF) target genes identified by ChIP-Atlas.

Data S6. Gene set enrichment analysis (GSEA) using Hallmark gene sets.

Data S7. Gene set enrichment analysis (GSEA) using TF target gene sets (Data S5).

## Materials and Methods

### Ethics Statement

This study was approved by the Review Boards of Johns Hopkins School of Medicine and Memorial Sloan Kettering Cancer Center.

### Patient selection

A cohort of 156 cases from the Gastrointestinal Cancer Rapid Medical Donation Program at Johns Hopkins Hospital and six cases from the Medical Donation Program at Memorial Sloan Kettering Cancer Center were used. All patients had a premortem diagnosis of PDAC based on pathologic review of resected or biopsy material and/or radiographic and biomarker studies. In addition, hematoxylin and eosin (H&E) stained sections of 30 resected PDACs were used for histologic review.

### Histology and Immunohistochemistry

H&E slides cut from all formalin-fixed and paraffin-embedded (FFPE) blocks of each autopsy were reviewed by two gastrointestinal pathologists (A.H. and C.I.D). Based on review and joint discussion a consensus diagnosis was rendered. Immunolabeling was performed on unstained serial sections cut from a subset of FFPE blocks per patient with antibodies against p63 (Ventana, clone 4A4) and CK5/6 (Ventana, clone D5/16B4) according to optimized protocol on a Ventana Benchmark XT autostainer (Ventana Medical Systems Inc.). Appropriate positive and negative controls were included in each run. Proportion of squamous differentiation in each carcinoma was estimated based on the number of blocks with SF/SD and the area of squamous differentiation within each block (1% tile for 1-5, 5% tile for 5-100%).

### Histologic Review for Entosis

All H&E sections of each case were reviewed for entotic cell-in-cell structures (CIC) using the criteria proposed by MacKay (*1*): cytoplasm of the host cell (winner or engulfing cell), nucleus of the host cell (typically crescent-shaped, binucleate, or multilobular and pushed against the cytoplasmic wall), an intervening vacuolar space completely surrounding the internalized cell (loser), cytoplasm of internalized cell, and nucleus of internalized cell (often round in shape and located centrally or acentrically). If internalized and/or engulfing cells were undergoing mitosis they were excluded from analysis. Any cases in which we were unable to count 50 high power fields and/or had less than five slides for review were excluded from this analysis. Representative entotic CICs were validated by immunofluorescence labeling for e-cadherin in combination with DAPI to highlight cell nuclei in the Molecular Cytogenetics Core at MSKCC (see MYC Immuno-FISH analysis section below for details).

### RNA Sequencing

Frozen sections were cut from samples for histologic review and regions of interest were macrodissected for extracting total RNA using TRIzol (Life Technologies) followed by Rneasy Plus Mini Kit (Qiagen). Each RNA sample was initially quantified by Qubit 2.0 Fluorometer (Thermo Fisher Scientific). Samples were additionally quantified by RiboGreen and assessed for quality control using an Agilent BioAnalyzer in the Integrated Genomics Core at MSKCC, and 513ng-1µg of total RNA with an RNA integrity number ranging from 1.3 to 8.3 underwent ribosomal depletion and library preparation using the TruSeq Stranded Total RNA LT Kit (Illumina catalog # RS-122-1202) according to instructions provided by the manufacturer with 8 cycles of PCR. Samples were barcoded and run on a HiSeq 4000 in a 100bp/100bp or 125/125bp paired end run, using the HiSeq 3000/4000 SBS Kit (Illumina). On average, 94 million paired reads were generated per sample and 26% of the data mapped to the transcriptome.

### RNA sequencing data alignment and analysis

Output data (FASTQ files) were mapped to the target genome using the rnaStar aligner (*2*) that maps reads genomically and resolves reads across splice junctions. We used the 2 pass mapping method outlined by Engstrom (*3*) in which the reads were mapped twice, the first mapping using a list of known annotated junctions from Ensembl and the second mapping based on known and novel junctions. Postprocessing of the output SAM files was performed using PICARD tools to add read groups and covert it to a compressed BAM format. The expression count matrix from the mapped reads was determined using HTSeq (www-huber.embl.de/users/anders/HTSeq) and the raw count matrix generated by HTSeq was processed using the R/Bioconductor package DESeq (www-huber.embl.de/users/anders/DESeq) to normalize the entire dataset between sample groups. Normalized log2 expression were used for downstream analyses (Supplementary Data Table 3).

### Expression type classification and PCA analysis

A 50 pancreatic cancer related gene set identified by Moffitt et al. was used to classify all samples into “classical” and “basal” types (*4*). Clustering analysis and heatmaps were displayed using the R package ‘pheatmap’ using spearman’s rank correlation. These 50 gene signatures were also used for generating the Primary Component Analysis (PCA) plot using DESeq2 package (https://www.bioconductor.org/packages/release/bioc/html/DESeq2.html).

### Gene set enrichment analysis

Gene set enrichment analysis was performed based on methods described (*5*). Both gene sets and transcription factor target gene sets (Supplementary Data Table 5) based on ChIP-seq data downloaded from ChIP-Atlas (http://chip-atlas.org/) (*6*) were used for analysis. Only TOP 500 ChIP peaks located within 1000bp from the TSS with scores over 50 were used.

### Network analysis and Cytoscape visualization

Co-expression networks were constructed by first identifying the best predicted soft threshold for transforming the data. Pearson correlation between any two genes across samples was next used as the weight between nodes. Subset of Keratins (KRT) family genes were used to construct the weighted gene-gene network, and the network structure was visualized using Cytoscape (*7*). We adjusted the width of edges connecting nodes based on the weights, and weights that are less than 0.05 were removed from the network.

### DNA sequencing

Genomic DNA was extracted from each tissue using QIAamp DNA Mini Kits (Qiagen). Whole genome sequencing (WGS) and whole exome sequencing (WES) and alignment performed as previously described (*8, 9*). Briefly, an Illumina HiSeq 2000 platform was used to target a coverage of 60X for WGS samples and 150X for WES samples. The resulting sequencing reads were analyzed in silico to assess quality, coverage, as well as alignment to the human reference genome (hg19) using BWA(*10*). After read de-duplication, base quality recalibration, and multiple sequence realignment were completed with the Picard Suite and GATK version 3.1(*11, 12*), somatic SNVs/INDELs were detected using Mutect version 1.1.6 and HaplotypeCaller version 2.4(*11, 13*). We excluded low-quality or poorly aligned reads from phylogenetic analysis. Filtering of called somatic mutations required each mutant to be observed in at least one neoplastic sample per patient with at least 5% variant allele frequency and with at least 20× coverage; correspondingly, each mutant must have been observed in less than 2% of the reads (or less than 2 reads total) of the matched normal sample with at least 10× coverage. Copy number analyses were performed using FACETS as previously described (*14*). Regarding PAM02, we used the data previously reported (*8*).

### Driver gene annotations

All somatic variants causing a frameshift deletion, frameshift insertion, in-frame deletion, in-frame insertion, non-synonymous missense, nonsense, nonstop, splice site/region, or a translation start site change were considered. Variants were called driver mutations if they passed at least three of the following methods: 20/20+ (*15*), 20/20+ PDAC(*15*), TUSON(*16*) and MutSigCV(*17*). For frameshift deletions, frameshift insertions and nonsense mutations specifically, passing only two of these four methods were required. Additionally, we required a CHASM p-value of ≤ 0.05 and an FDR of ≤ 0.25 for the 20/20+ and 20/20+ PDAC methods. We also considered genes significantly mutated in large PDAC sequencing studies (*18–20*). All driver gene alternation was also manually reviewed with Integrative Genome Viewer (*21*).

### Whole Genome Duplication

Whole genome duplication (WGD) was performed in combination of computational analysis and manually reviewed following Bielski et al.(*22*), called if MCN ≥ 2 and ploidy ≥ 2.5, and >50% of the autosomal genome was affected. Three low tumor purity samples (PAM22PT5, PAM25PT2 and PAM32PT4) which didn’t match these criteria were judged in consideration of expecting WGD occurrent point in phylogenetic trees.

### Evolutionary analysis

We derived phylogenies for each set of samples by using Treeomics 1.7.9 (*23*). Each phylogeny was rooted at the matched patient’s normal sample and the leaves represented tumor samples. Treeomics employs a Bayesian inference model to account for error-prone sequencing and varying neoplastic cell content to calculate the probability that a specific variant is present or absent. The global optimal tree is based on Mixed Integer Linear programming. All evolutionary analyses were performed based on WES data with exception of PAM02 (WGS and additional target sequencing)(*8*) and MPAM06 (WGS). Somatic alterations present in all analyzed samples of a PDAC were considered truncal, in a subset of samples considered branched, and in a single sample considered private.

### *MYC* amplification

*MYC* amplification was defined as ≥6-fold by FACETS (*14*) or FISH (see following chapter). In brief, FACETS performs a complete analysis that includes library size and GC-normalization, and segmentation of total and allele-specific signals, using coverage and genotypes of single nucleotide polymorphisms simultaneously across the exome. The resulting segments accurately identify points of change in the exome, accounting for diploidy, purity, and average ploidy for each sample. A maximum likelihood approach then assigns each segment with a major and minor integer copy number.

### *MYC* Immuno-FISH analysis

Immuno-FISH was performed on paraffin sections according to procedures optimized at the Molecular Cytogenetics Core Facility. The primary (E-Cadherin [24E10] Rabbit mAB) and secondary (Goat anti-Rabbit Alexa 488) antibody was purchased form Cell Signaling Technology and Invitrogen (Thermo Fisher Scientific) respectively. The 2-color *MYC*/Cen8 probe was prepared in-house and consisted of BAC clones containing the full length *MYC* gene (clones RPI-80K22, RP11-1136L8, and CTD-2267H22; labeled with Red dUTP) and a centromeric repeat plasmid for chromosome 8 served as the control (pJM128; labeled with Green dUTP). Briefly, de-waxed paraffin sections were microwaved in 10mM sodium citrate, pretreated with 10% pepsin for 10 minutes at 37oC, rinsed in 2XSSC, dehydrated in ethanol series (70%, 90% and 100%), co-denatured at 80oC for 4 minutes with 5-20uL of *MYC*/Cen8 DNA-FISH probe, and hybridized for 72 hours at 37oC. Following hybridization, sections were washed with wash buffer (0.01% Tween 20 in 2XSSC), fixed in 4% formaldehyde for 15-20 minutes at RT, rinsed in 1XPBS, blocked at RT for 1 hour (blocking buffer: 5% FBS and 0.01% Tween 20 in 1XPBS), and incubated overnight at 4oC with primary antibody (1:100)(dilution buffer: 1% FBS and 0.01% Tween 20 in 1XPBS). Following overnight incubation, sections were washed with wash buffer, rinsed in 1XPBS, incubated with secondary antibody (1:500) for 1 hour at 37oC, rinsed in 1XPBS, stained with DAPI and mounted in antifade (Vectashield, Vector Laboratories). Slides were scanned using a Zeiss Axioplan 2i epifluorescence microscope equipped with Isis 5.5.9 imaging software (MetaSystems Group Inc, Waltham, MA). Metafer and VSlide modules within the software were used to generate virtual image of H&E and DAPI-stained sections. In all, corresponding H&E sections assisted in localizing tumor region and histology (GL, SF or SD). The entire section was systematically scanned under 63 × objectives to assess *MYC*/Cen8 copy number across different histologies and to identify entotic cell-in-cell structures (CIC). All observed entotic cells and representative regions within a case were imaged through the depth of the tissue (merged stack of 16 z-section images taken at 0.5 micron intervals) and signal counts performed on captured images. For correlation of *MYC*/Cen8 copy number with histology, for each case, a minimum of 50 discrete nuclei were scored (range 50-150). Within a given histology (GL, SF or SD), when *MYC*/Cen8 copy number was heterogenous and topographically distinct, a minimum of 50 discrete nuclei were scored for each distinct region whenever possible. For correlation of *MYC*/Cen8 copy number with entosis, only CICs meeting the selection criteria previously described were scored. For each CIC, *MYC*/Cen8 copy number was recorded separately for the “winner” and “loser”. Presence of E-Cadherin staining (which highlights the cell perimeter) and nuclear morphology helped distinguish the “loser” (internalized cell with uniformly round nucleus) from “winner” (host cell with crescent-shaped, binucleate, or multilobulated nucleus and often pushed against the cytoplasmic wall). To minimize truncation artifacts, only nuclei with at least 1 signal for *MYC* and Cen8 were selected. *MYC* amplification was defined as: ≥2 *MYC*/Cen8 ratio, ≥6 copies of *MYC* (discrete signal) or presence of at least one *MYC* cluster (≥4 copies; tandem duplications). 3~5 copies of *MYC*/Cen8 were regarded as copy number gain (polysomy).

### Statistics

All statistics was performed using XLSTAT (version 2018.2).

---

**Fig. S1:**
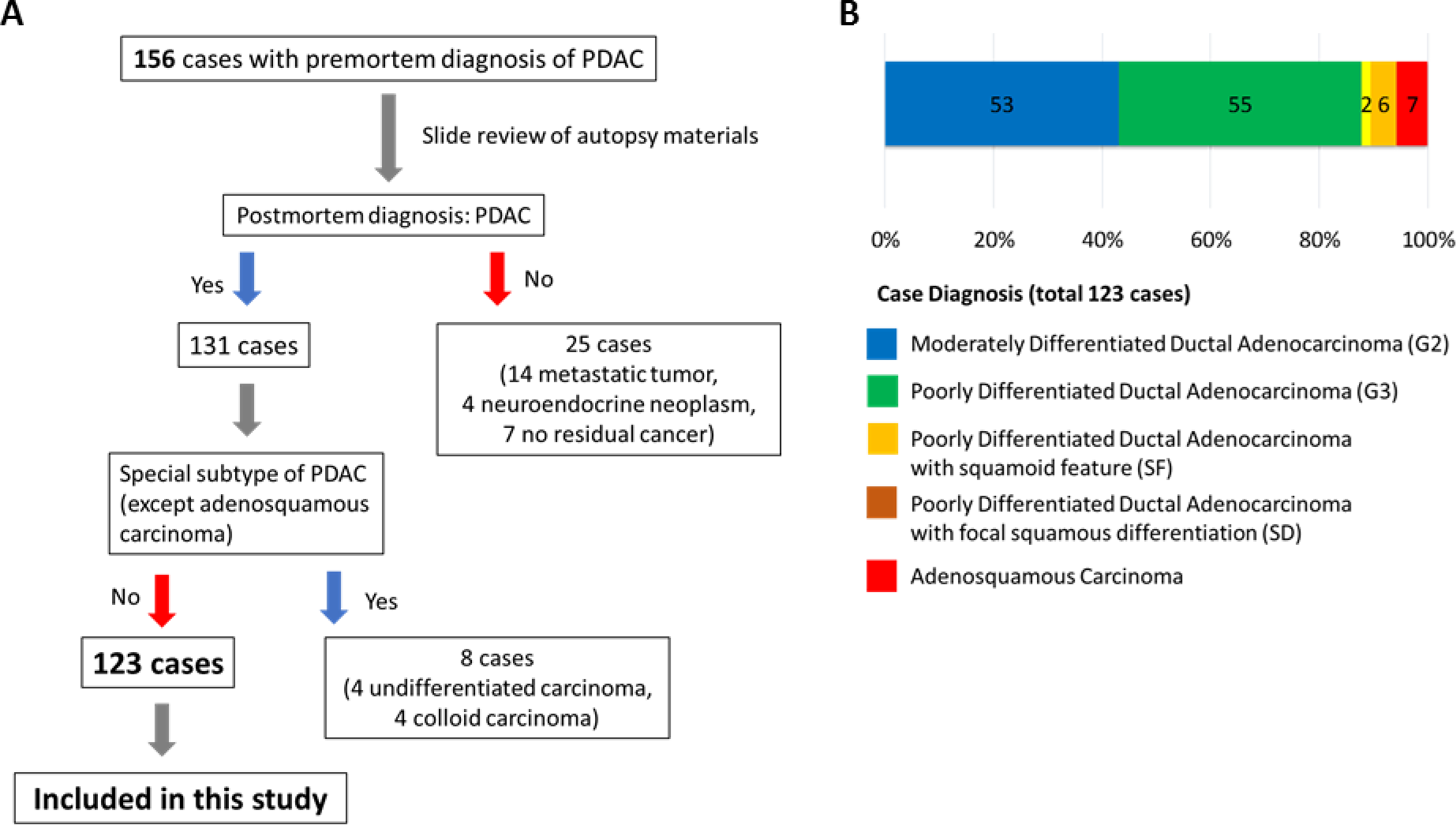
Case selection and postmortem diagnosis. (A) Schematic of case selection for current study. (B) Postmortem case diagnoses. Seven cases corresponded to adenosquamous carcinoma (ASC), four cases showed focal (<30%) squamous differentiation (SD), and two cases showed focal squamoid features (SF).

**Fig. S2:**
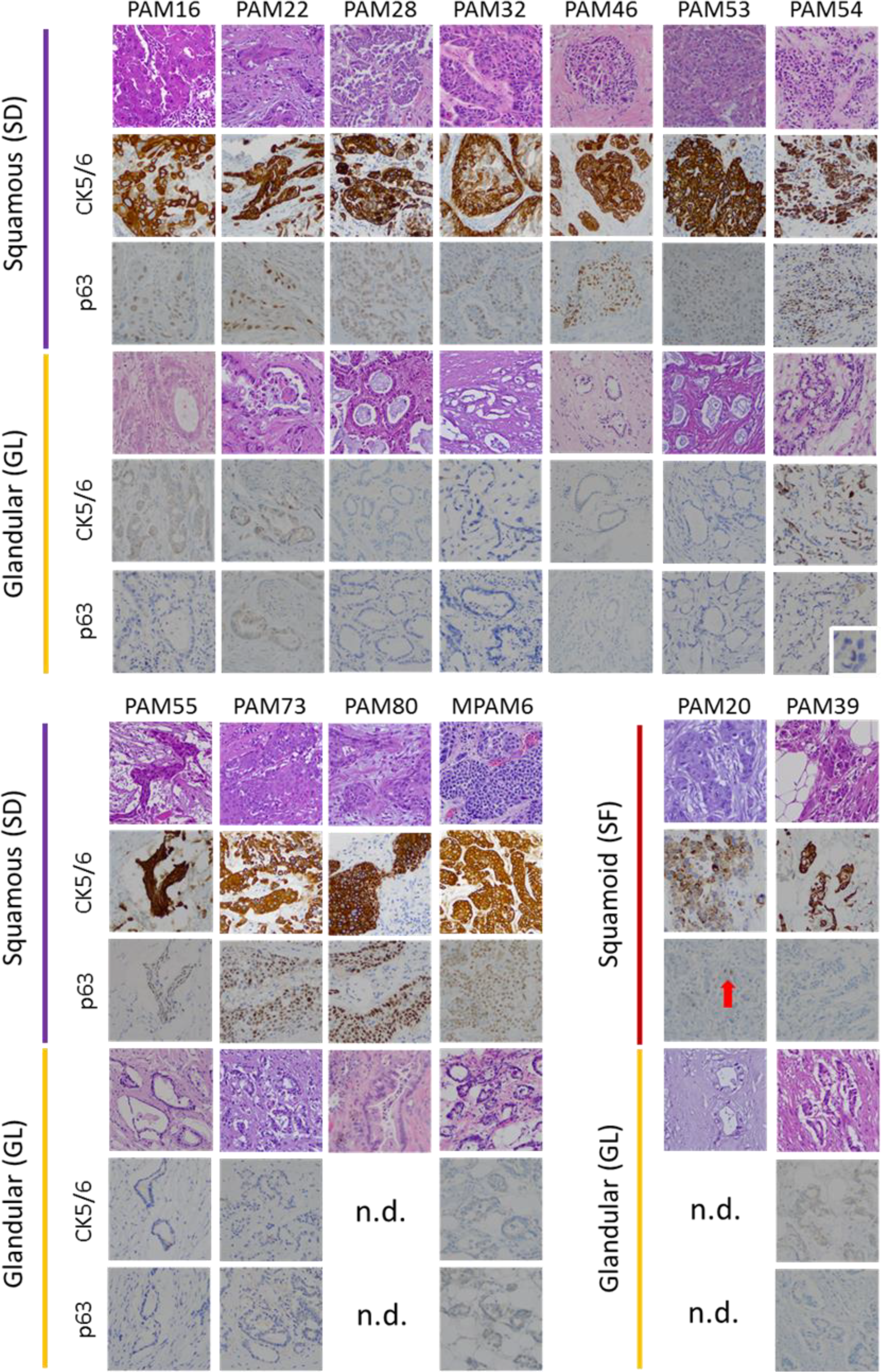
Immunolabelling for glandular and squamous components in 13 Representative PDACs. All regions with squamous differentiation (SD) showed positivity for CK5/6 and p63, whereas no labeling was observed in regions with glandular morphology (GL). In two PDACs the neoplastic cells stained positive for CK5/6 but were negative for p63 and thus classified as having squamoid features. n.d., no immunolabeling performed.

**Fig. S3:**
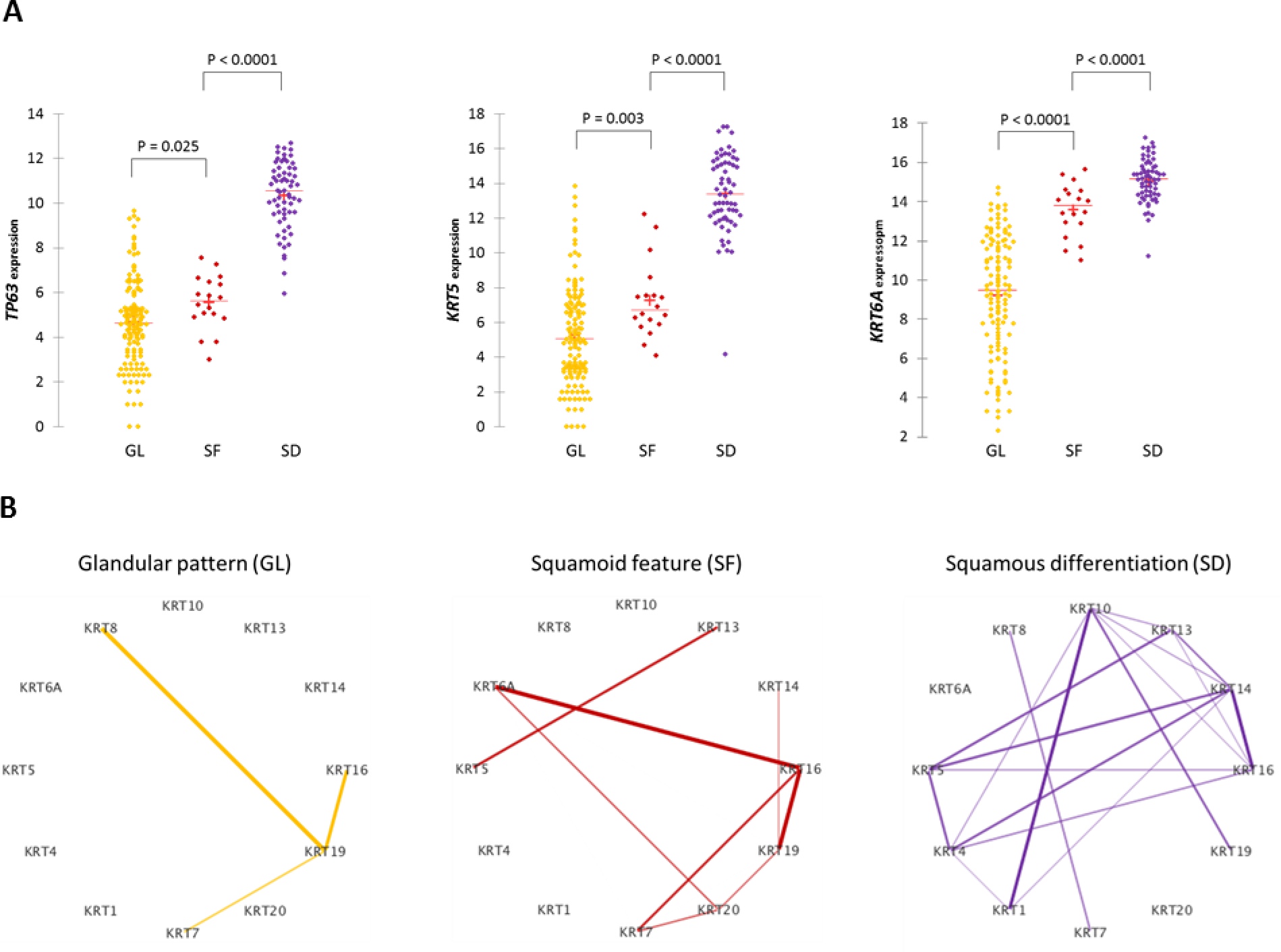
mRNA Expression of squamous markers in samples with glandular growth pattern (GL), squamoid features (SF) and squamous differentiation (SD). **a**. mRNA expression of *TP63, KRT5* and *KRT6A*. SD have higher expression *of TP63, KRT5* and *KRT6A* than GL. SF have intermediate expression pattern between SD and GL. b. KRT network based on mRNA expression. In GL, *KRT19* (normally expressed in ductal epithelia) is a hub in pancreas cancer. In SF, *KRT6A* and *KTR5* (normally expressed in squamous epithelium) have some interaction. In SD, stratified squamous epithelium keratins (*KRT4, KRT5, KRT14, KRT15*) and heavy weight keratins (*KRT*1 and *KRT1*0) were expressed in the network.

**Fig. S4.**
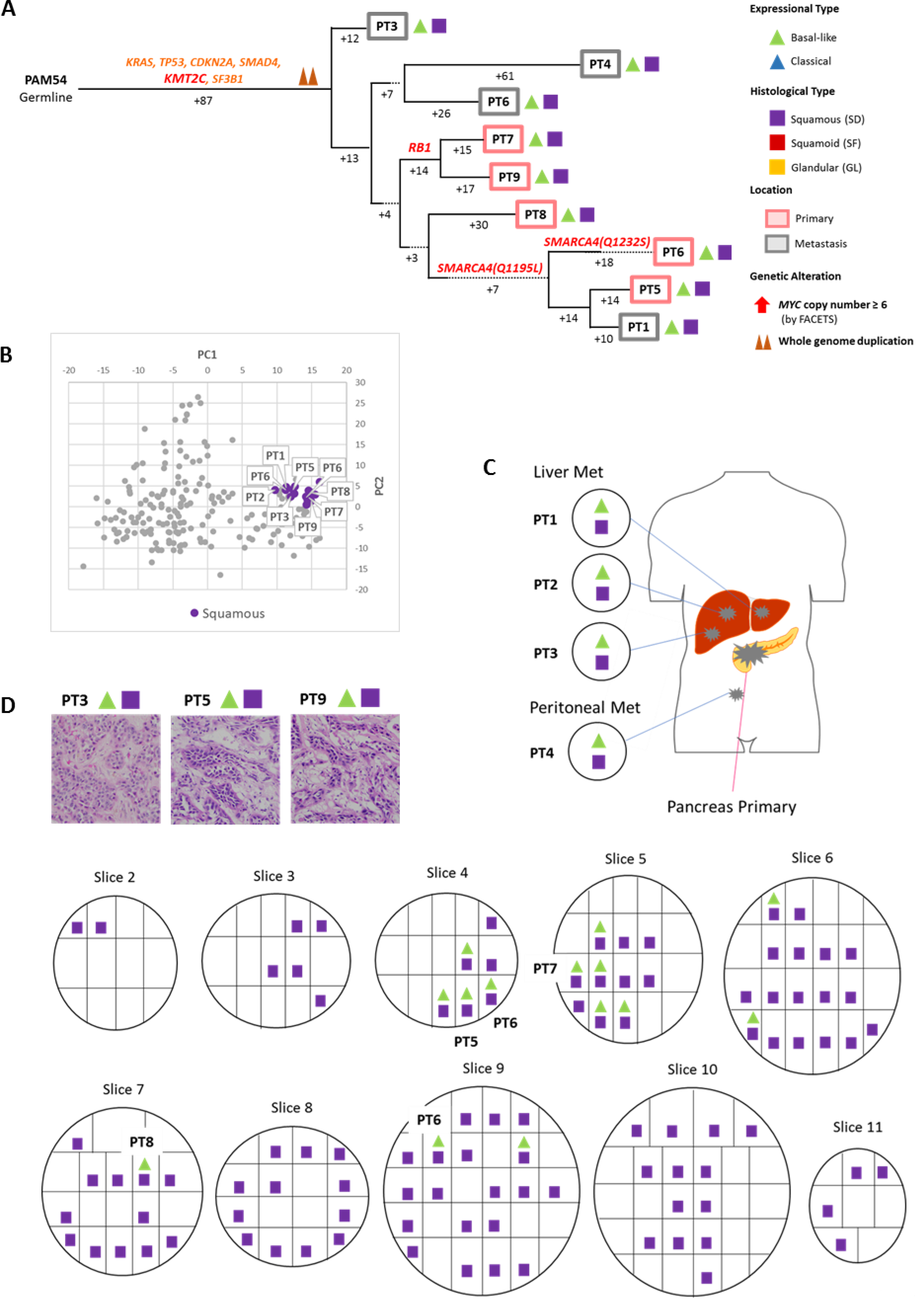
Integration of Transcriptomic and Morphologic Features with Phylogenetic Patterns in Pancreatic Ductal Adenocarcinoma PAM54 with Truncal *KMT2C* mutation. **(A)** Phylogenetic analysis illustrating the clonal relationship of samples analyzed in this patient. The predicted timing of somatic alterations in driver genes and whole genome duplication are also shown. Mutations in chromatin modifier genes are in red font, all others in orange. Truncal driver genes are notable for a *KMT2C* somatic alteration, whereas mutations in *RB1* and *SMARCA4* (two independent mutations) are present in a subset of samples. SD in this carcinoma was found in all samples analyzed, although it was admixed with a minor GL component in some samples. **(B)** Principal components analysis illustrating highly similar expression between samples. **(C)** Relationship of anatomic location to morphologic and transcriptional profiles. **(D)** Representative histologic images of tumors in the same patient.

**Fig. S5.**
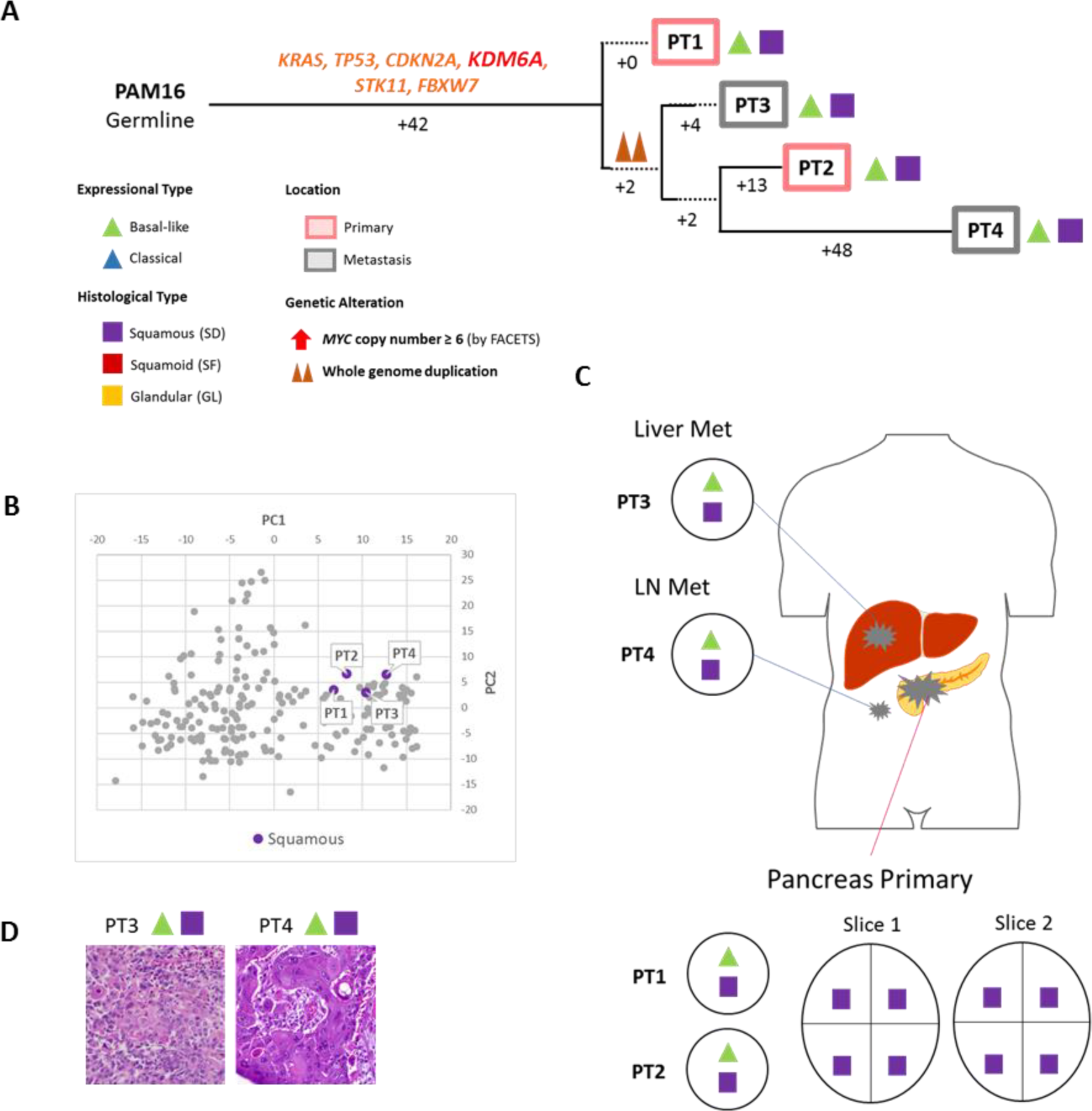
Integration of Transcriptomic and Morphologic Features with Phylogenetic Patterns in Pancreatic Ductal Adenocarcinoma PAM16 with Truncal *KDM6A* mutation. **(A)** Phylogenetic analysis illustrating the clonal relationship of samples analyzed in this patient. The predicted timing of somatic alterations in driver genes and whole genome duplication are shown. Mutations in chromatin modifier genes are in red font, all others in orange. Truncal driver genes are notable for a *KDM6A* somatic alteration. SD in this carcinoma was found in all samples analyzed, although it was admixed with a minor GL component in some samples. **(B)** Principal components analysis illustrating highly similar expression between samples. **(C)** Relationship of anatomic location to morphologic and transcriptional profiles. **(D)** Representative histologic images of metastatic tumors PT3 and PT4.

**Fig. S6.**
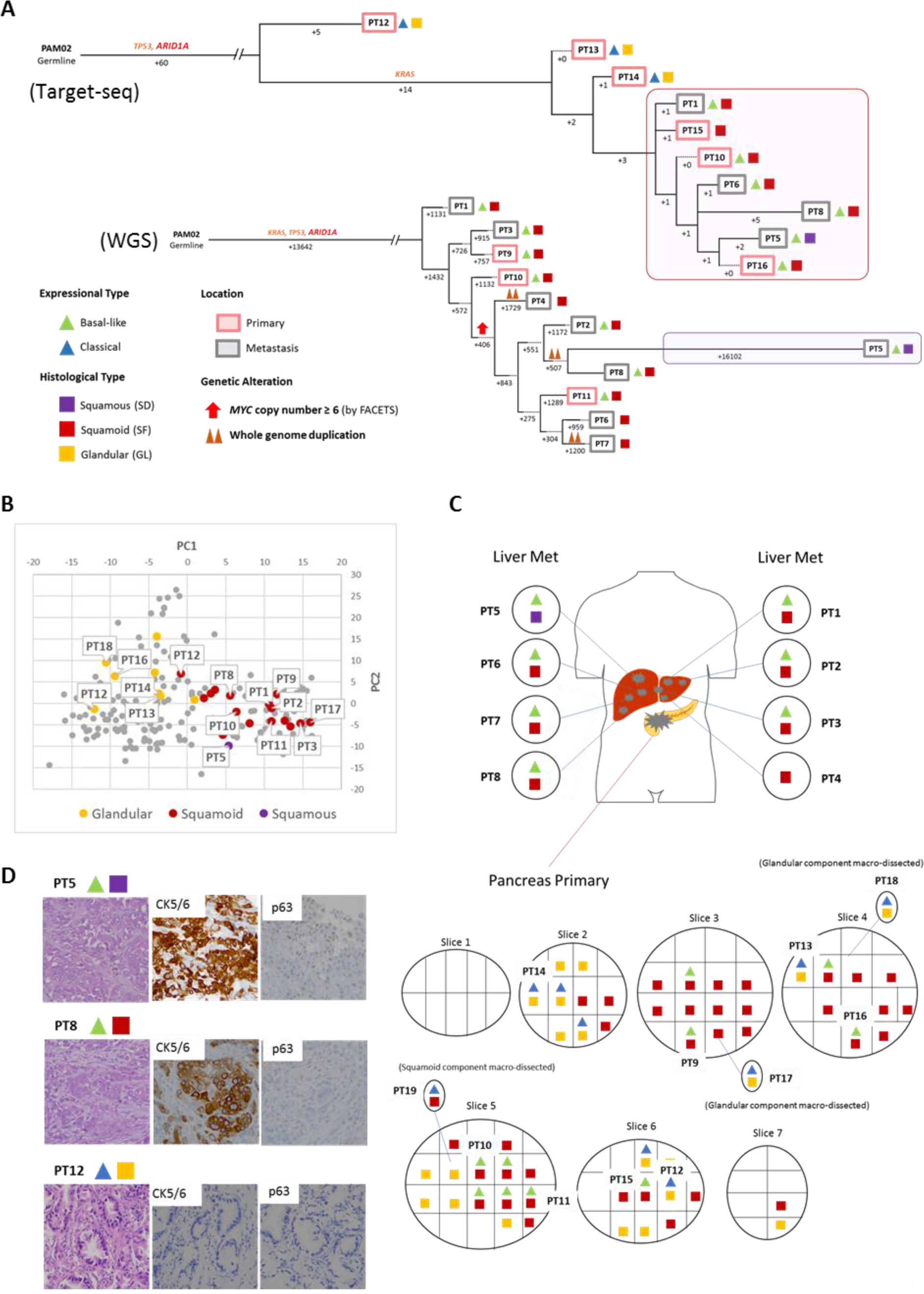
Integration of Transcriptomic and Morphologic Features with Phylogenetic Patterns in Pancreatic Ductal Adenocarcinoma PAM02 with Truncal *ARID1A* mutation. **(A)** Phylogenetic analysis illustrating the clonal relationship of samples analyzed in this patient. The predicted timing of somatic alterations in driver genes, whole genome duplication and *MYC* amplification are shown. Mutations in chromatin modifier genes are in red font, all others in orange. Red and purple outlines indicate samples that have SF/SD based on RNAseq (triangles) and/or histology (squares). Phylogenetic analysis based on WGS (top) or targeted sequencing (bottom) of an overlapping set of samples from this patient. Truncal driver genes are notable for an *ARID1A* somatic alteration. Primary and metastatic samples with SF/SD in this patient are clonally related. **(B)** Principal components analysis illustrating the divergent expression profiles between GL and SF/SD morphologies present in the primary tumor. **(C)** Relationship of anatomic location to morphologic and transcriptional heterogeneity. Both GL and SF were seen in the primary tumor with corresponding “classic” or “basal-like” expression profiles respectively. One liver metastasis (PT5) showed SD. **(D)** Representative histologic and immunohistochemical images of metastasis samples PT5 and PT8 and primary tumor samples PT12.

**Fig. S7.**
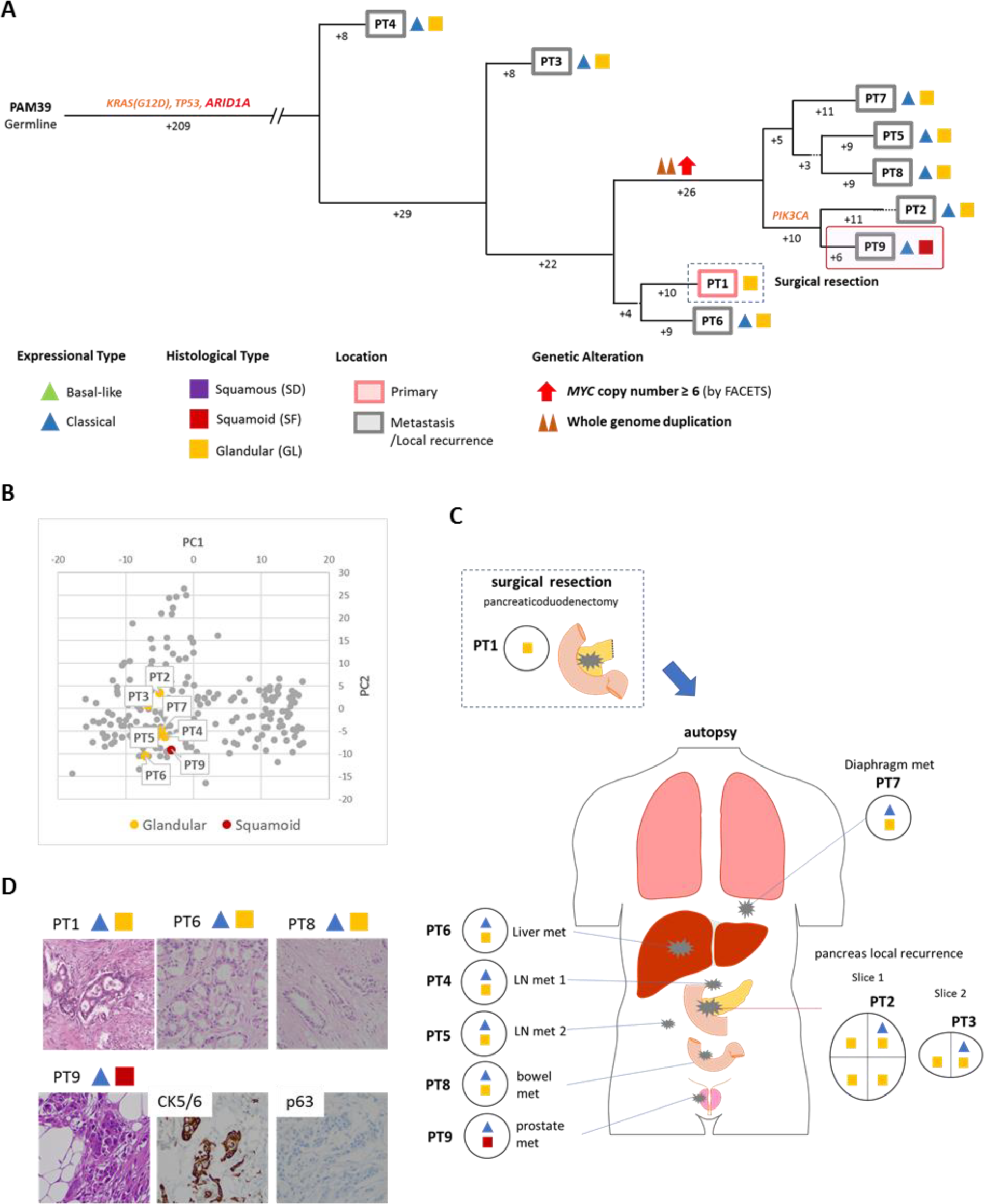
Integration of Transcriptomic and Morphologic Features with Phylogenetic Patterns in Pancreatic Ductal Adenocarcinoma PAM39 with Truncal *ARID1A* mutation. **(A)** Phylogenetic analysis illustrating the clonal relationship of samples analyzed in this patient. The predicted timing of somatic alterations in driver genes, whole genome duplication and *MYC* amplification are shown. Mutations in chromatin modifier genes are in red font, all others in orange. Red outline indicates the one sample with SF based on histology and immunohistochemical analysis (squares) but a classic-type expression profile (triangle). Truncal driver genes are notable for an *ARID1A* somatic alteration. SF is confined to one prostate metastasis sample (PT9). **(B)** Principal components analysis shows a similar gene expression profile between the samples with GL or SF morphology. **(C)** Relationship of anatomic location to morphologic and transcriptional heterogeneity. (**D)** Representative histologic and immunohistochemical images of the primary tumor (PT1) and prostate metastasis (PT9).

**Fig. S8.**
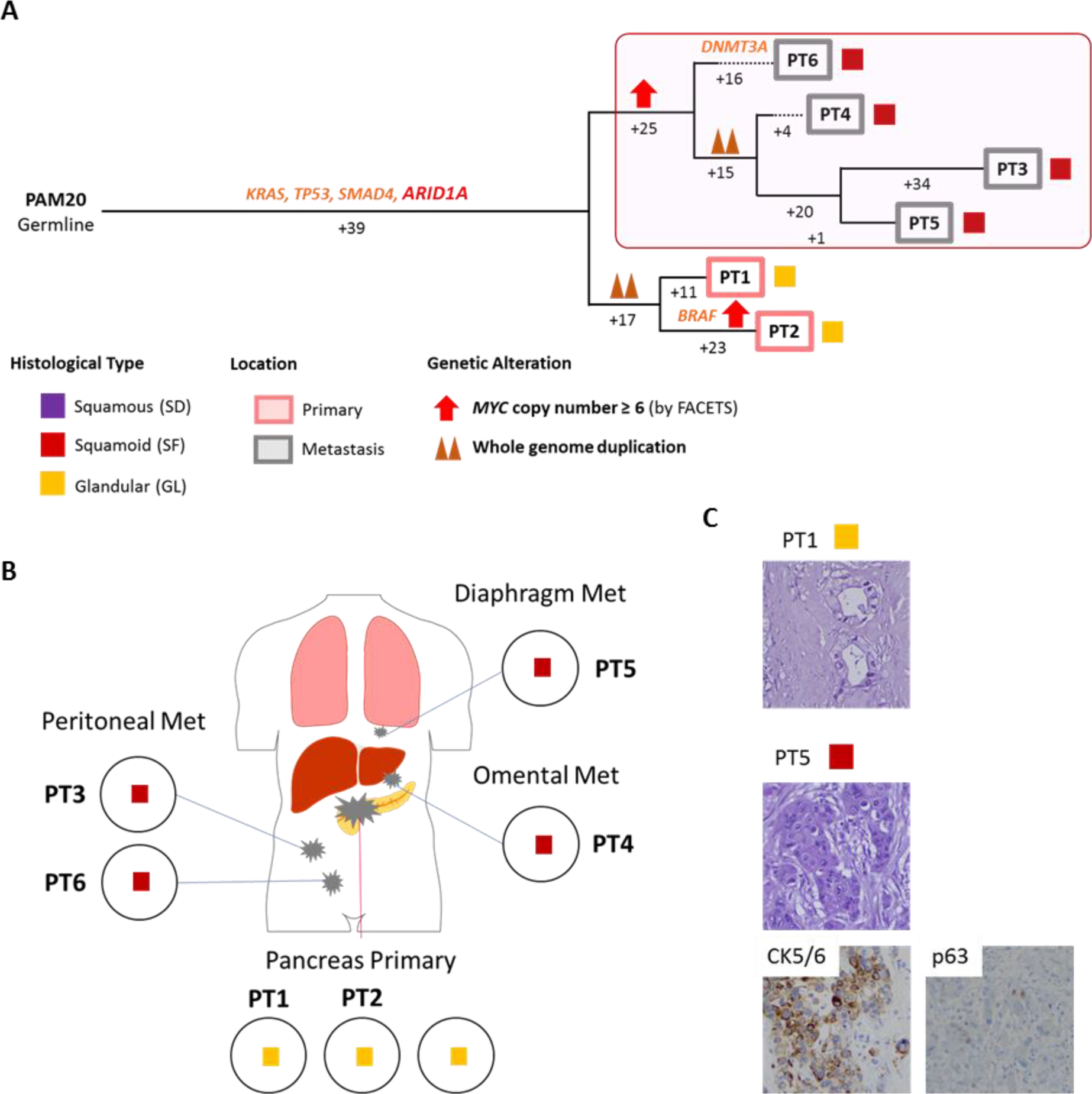
Integration of Transcriptomic and Morphologic Features with Phylogenetic Patterns in Pancreatic Ductal Adenocarcinoma PAM20 with Truncal *ARID1A* mutation. **(A)** Phylogenetic analysis illustrating the clonal relationship of samples analyzed in this patient. The predicted timing of somatic alterations in driver genes, whole genome duplication and *MYC* amplification are shown. Mutations in chromatin modifier genes are in red font, all others in orange. Red outline indicates the samples that have SF based on histology and immunolabeling (squares). Truncal driver genes are notable for a *ARID1A* somatic alteration. *MYC* amplification (>6 copies) was detected in all samples with SF, and in a phylogenetically distinct sample with GL within the primary tumor. Samples with SF in this carcinoma (PT3-PT6) are clonally related. (B) Relationship of anatomic location to morphologic heterogeneity. The metastasis samples PT3-PT6 showed SF whereas the primary tumor samples showed GL. **(C)** Representative histologic and/or immunohistochemical images of the primary tumor (PT1) and diaphragm metastasis (PT5).

**Fig. S9.**
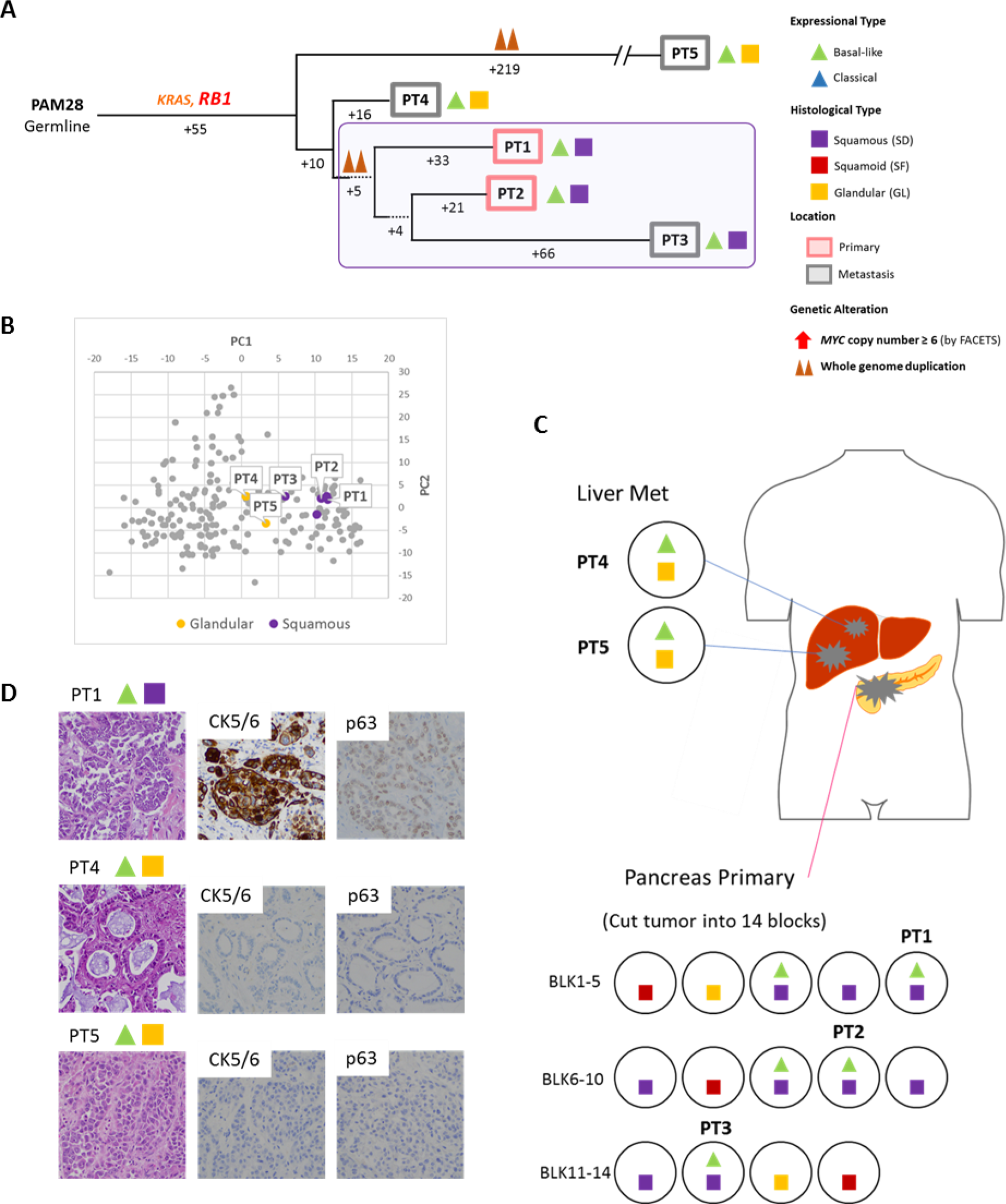
Integration of Transcriptomic and Morphologic Features with Phylogenetic Patterns in Pancreatic Ductal Adenocarcinoma PAM28 with Truncal *RB1* Mutation. **(A)** Phylogenetic analysis illustrating the clonal relationship of samples analyzed in this patient. The predicted timing of somatic alterations in driver genes and whole genome duplication are shown. The mutation in *RB1* is in red font, all others in orange. Purple outline indicates samples that have SD based on RNAseq (triangles) and/or histology (squares). Truncal driver genes are notable for an *RB1* somatic alteration. Samples with SD are more related to each other than to other samples in this patient. **(B)** Principal components analysis shows that samples PT1-PT3 with “basal-like”expression and SD morphology are distinct from samples PT4 and PT5 that have “basal-like” expression but GL morphology. **(C)** Relationship of anatomic location to morphologic and transcriptional heterogeneity. Both GL and SF were seen in the primary tumor, yet liver metastases PT4 and PT5 have GL morphology and a “basal-type”expression profile. **(D)** Representative histologic images and immunohistochemical labeling of primary tumor sample PT1 and liver metastases PT4 and PT5.

**Fig. S10.**
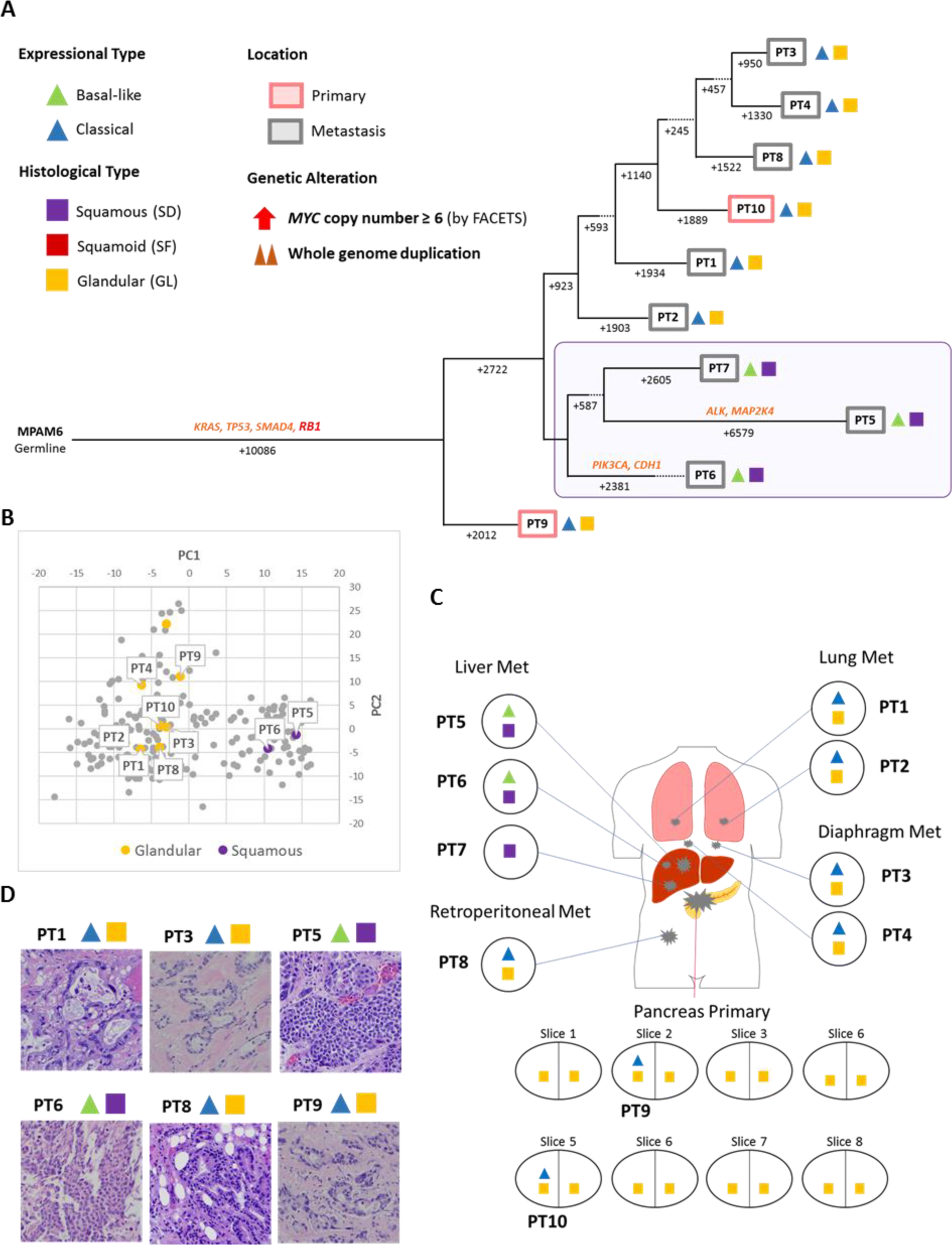
Integration of Transcriptomic and Morphologic Features with Phylogenetic Patterns in Pancreatic Ductal Adenocarcinoma MPAM6 with deleterious truncal *RB1* mutation. **(A)** Phylogenetic analysis illustrating the clonal relationship of samples analyzed in this patient. The predicted timing of somatic alterations in driver genes are shown. Mutations in *RB1* are in red font, all others in orange. Purple outline indicates samples that have SD based on RNAseq (triangles) and/or histology (squares). Truncal driver genes are notable for a deleterious *RB1* mutation. Samples with SD (PT5-PT7) are more related to each other than to other samples in the same patient. **(B)** Principal components analysis indicates distinct gene expression profiles between GL and SF samples. **(C)** Relationship of anatomic location to morphologic and transcriptional heterogeneity. SF is confined to the liver metastases (PT5-PT7). **(D)** Representative histologic images of the primary and multiple metastatic tumors in the same patient.

**Fig. S11.**
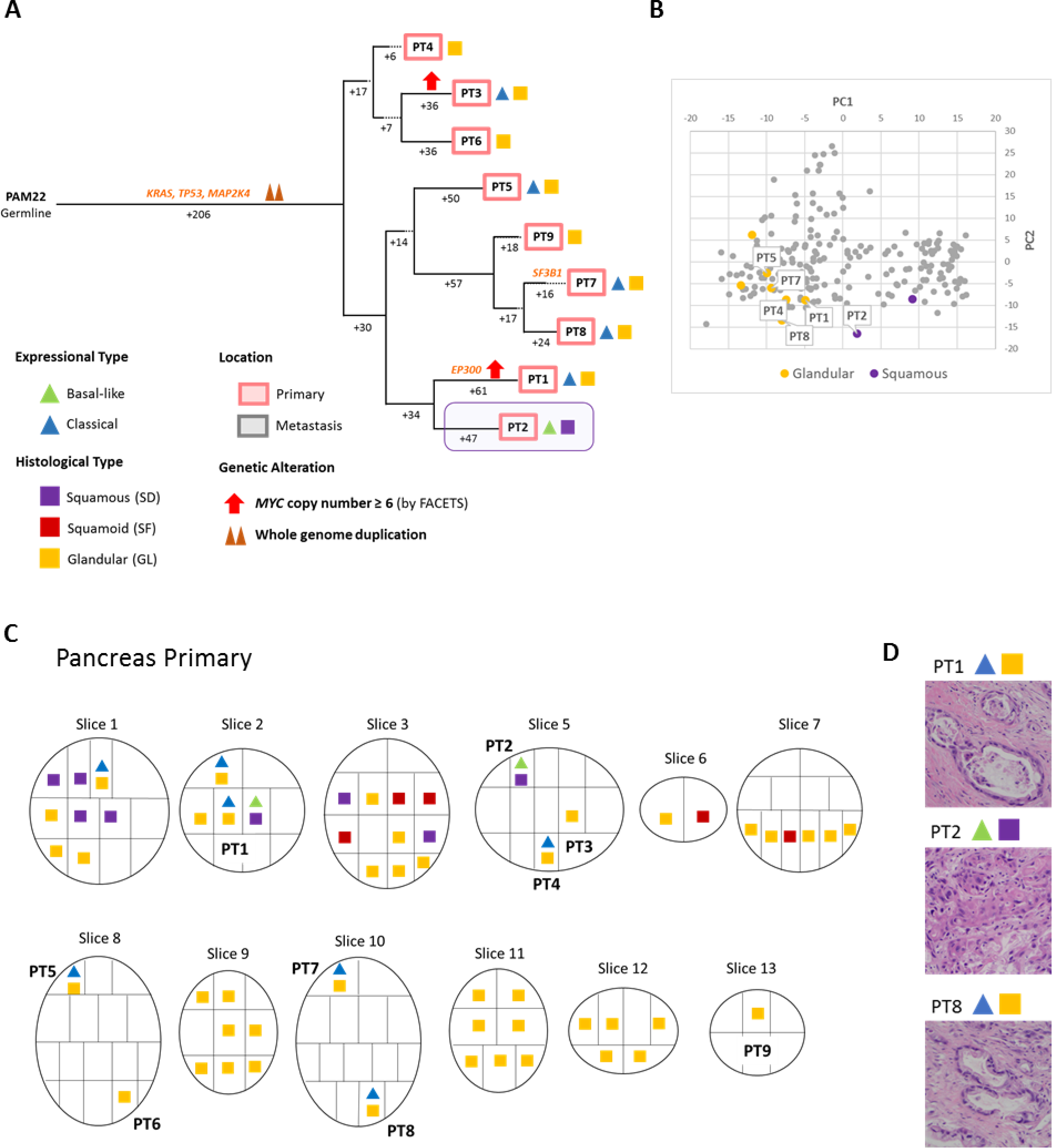
Integration of Transcriptomic and Morphologic Features with Phylogenetic Patterns in Pancreatic Ductal Adenocarcinoma PAM22. **(A)** Phylogenetic analysis illustrating the clonal relationship of samples analyzed in this patient. Purple outline indicates samples that have SD based on RNAseq (triangles) and histology/immunohistochemistry (squares). The predicted timing of somatic alterations in driver genes, whole genome duplication and *MYC* amplification are also shown. SD is confined to a single sample within the multiregion sampled primary tumor (PT2). **(B)** Principal components analysis indicates that PT2 shows a different expression profile from all other primary tumor samples that have GL morphology. **(C)** Relationship of anatomic location within the primary tumor to morphologic and/or transcriptional heterogeneity for SF/SD. **(D)** Representative histologic images of representative tumors in the same patient.

**Fig. S12.**
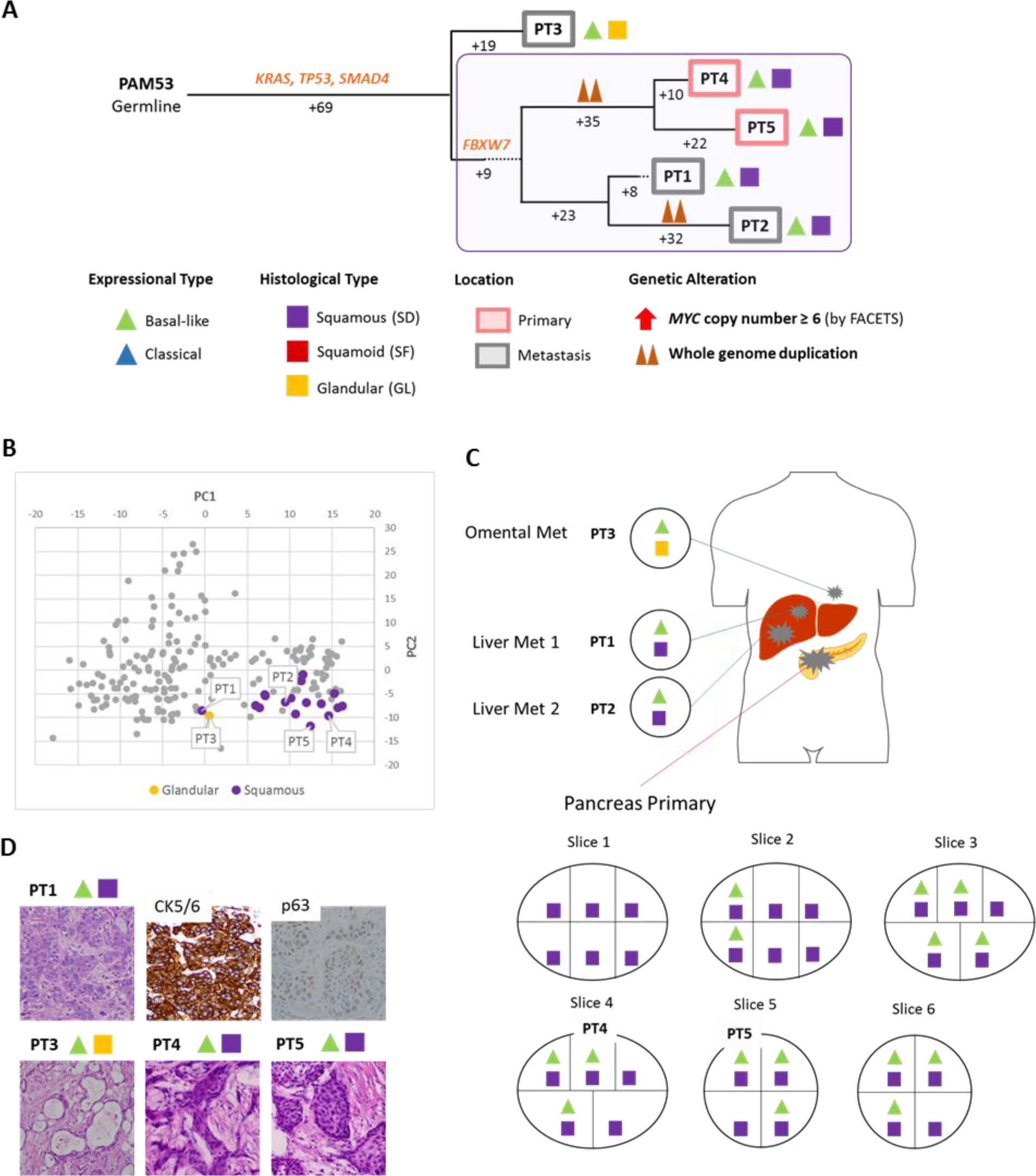
Integration of Transcriptomic and Morphologic Features with Phylogenetic Patterns in Pancreatic Ductal Adenocarcinoma PAM53. **(A)** Phylogenetic analysis illustrating the clonal relationship of samples analyzed in this patient. The predicted timing of somatic alterations in driver genes and whole genome duplication are also shown. Purple outline indicates samples that have SD based on RNAseq (triangles) and histology/immunohistochemistry (squares). The one sample with a classical expression profile and GL morphology forms the outgroup in the tree. Four samples with basal-like expression and SD correspond to both the primary tumor (PT4 and PT5) and metastasis (PT1 and PT2). **(B)** Principal components analysis indicates samples PT1 and PT3 have relatively different expression profiles from other SD samples. **(C)** Relationship of anatomic location to morphologic and transcriptional heterogeneity. SD was found in one omental metastasis (PT3) which is also showed “basal-like” expression. **(D)** Representative histologic and immunohistochemical images of the primary tumor samples PT4 and PT5, liver metastasis PT1 and omental metastasis PT3.

**Fig. S13.**
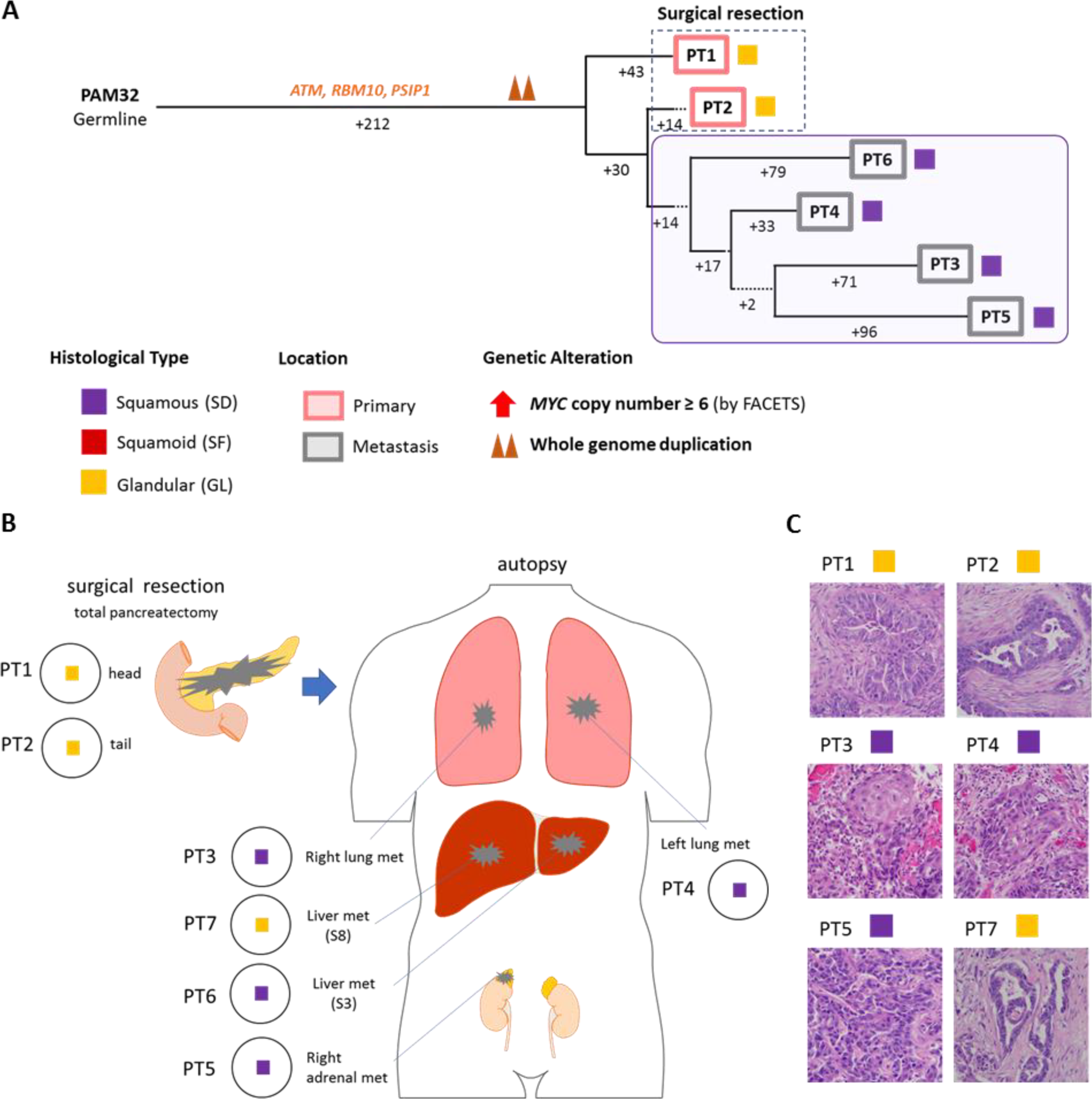
Integration of Transcriptomic and Morphologic Features with Phylogenetic Patterns in Pancreatic Ductal Adenocarcinoma PAM32. **(A)** Phylogenetic analysis illustrating the clonal relationship of samples analyzed in this patient. The predicted timing of somatic alterations in driver genes and whole genome duplication are also shown. Purple outline indicates samples that have SD based on histology and immunolabeling (squares). Samples PT3-PT6 with SD are more closely related to each other than to other samples in the same patient. **(B)** Relationship of anatomic location to morphologic heterogeneity. One liver metastasis (PT7, not sequenced) showed GL morphology. **(C)** Representative histologic images of the primary and metastatic tumors in this patient.

**Data S1. (separate Excel file)**

Clinical information of PDA with squamous differentiation or squamoid feature.

**Data S2. (separate Excel file)**

Sample information of RNA-seq, WES, WGS and target-seq.

**Data S3.** RNA expression, released after publication

**Data S4. (separate Excel file)**

Gene alternation of our cohort and MSK IMPACT Clinical Sequencing Cohort.

**Data S5. (separate Excel file)**

Transcription factor (TF) target genes identified by ChIP-Atlas.

**Data S6. (separate Excel file)**

Gene set enrichment analysis (GSEA) using Hallmark gene sets.

**Data S7. (separate Excel file)**

Gene set enrichment analysis (GSEA) using TF target gene sets (Data S5).

## References

1. T. Kamisawa, L. D. Wood, T. Itoi, K. Takaori, Pancreatic cancer. Lancet 388, 73–85 (2016).

2. S. Gillen, T. Schuster, C. Meyer Zum Buschenfelde, H. Friess, J. Kleeff, Preoperative/neoadjuvant therapy in pancreatic cancer: a systematic review and meta-analysis of response and resection percentages. PLoS Med 7, e1000267 (2010).

3. R. L. Siegel, K. D. Miller, A. Jemal, Cancer statistics, 2019. CA: a cancer journal for clinicians, (2019).

4. A. V. Biankin et al., Pancreatic cancer genomes reveal aberrations in axon guidance pathway genes. Nature 491, 399–405 (2012).

5. N. Waddell et al., Whole genomes redefine the mutational landscape of pancreatic cancer. Nature 518, 495–501 (2015).

6. P. Bailey et al., Genomic analyses identify molecular subtypes of pancreatic cancer. Nature 531, 47–52 (2016).

7. Cancer_Genome_Atlas_Research_Network, Integrated Genomic Characterization of Pancreatic Ductal Adenocarcinoma. Cancer Cell 32, 185–203 e113 (2017).

8. A. K. Witkiewicz et al., Whole-exome sequencing of pancreatic cancer defines genetic diversity and therapeutic targets. Nature communications 6, 6744 (2015).

9. R. E. Wilentz et al., Genetic, immunohistochemical, and clinical features of medullary carcinoma of the pancreas: A newly described and characterized entity. The American journal of pathology 156, 1641–1651 (2000).

10. E. A. Collisson et al., Subtypes of pancreatic ductal adenocarcinoma and their differing responses to therapy. Nature medicine 17, 500–503 (2011).

11. R. A. Moffitt et al., Virtual microdissection identifies distinct tumor-and stroma-specific subtypes of pancreatic ductal adenocarcinoma. Nature genetics 47, 1168–1178 (2015).

12. O. G. McDonald et al., Epigenomic reprogramming during pancreatic cancer progression links anabolic glucose metabolism to distant metastasis. Nature genetics 49, 367–376 (2017).

13. J. R. Brody et al., Adenosquamous carcinoma of the pancreas harbors KRAS2, DPC4 and TP53 molecular alterations similar to pancreatic ductal adenocarcinoma. Mod Pathol 22, 651–659 (2009).

14. D. E. Kardon, L. D. Thompson, R. M. Przygodzki, C. S. Heffess, Adenosquamous carcinoma of the pancreas: a clinicopathologic series of 25 cases. Mod Pathol 14, 443–451 (2001).

15. N. Fukushima et al., Ductal adnocarcinoma variants and mixed neoplasms of the pancreas. World Health Organization Classification of Tumors 4th Edition, 292–296 (2010).

16. C. A. Boyd, J. Benarroch-Gampel, K. M. Sheffield, C. D. Cooksley, T. S. Riall, 415 patients with adenosquamous carcinoma of the pancreas: a population-based analysis of prognosis and survival. J Surg Res 174, 12–19 (2012).

17. M. Overholtzer et al., A nonapoptotic cell death process, entosis, that occurs by cell-in-cell invasion. Cell 131, 966–979 (2007).

18. H. L. Mackay et al., Genomic instability in mutant p53 cancer cells upon entotic engulfment. Nature communications 9, 3070 (2018).

19. C. Liu et al., The UPF1 RNA surveillance gene is commonly mutated in pancreatic adenosquamous carcinoma. Nature medicine 20, 596–598 (2014).

20. J. Andricovich et al., Loss of KDM6A Activates Super-Enhancers to Induce Gender-Specific Squamous-like Pancreatic Cancer and Confers Sensitivity to BET Inhibitors. Cancer Cell 33, 512–526.e518 (2018).

21. A. Zehir et al., Mutational landscape of metastatic cancer revealed from prospective clinical sequencing of 10,000 patients. Nature medicine 23, 703–713 (2017).

22. I. Gonzalez-Vasconcellos et al., The Rb1 tumour suppressor gene modifies telomeric chromatin architecture by regulating TERRA expression. Scientific reports 7, 42056 (2017).

23. G. Lomberk et al., Distinct epigenetic landscapes underlie the pathobiology of pancreatic cancer subtypes. Nature communications 9, 1978 (2018).

24. C. Schleger, C. Verbeke, R. Hildenbrand, H. Zentgraf, U. Bleyl, c-MYC activation in primary and metastatic ductal adenocarcinoma of the pancreas: incidence, mechanisms, and clinical significance. Mod Pathol 15, 462–469 (2002).

25. M. Wirth, S. Mahboobi, O. H. Kramer, G. Schneider, Concepts to Target MYC in Pancreatic Cancer. Mol Cancer Ther 15, 1792–1798 (2016).

26. C. M. Bielski et al., Genome doubling shapes the evolution and prognosis of advanced cancers. Nature genetics 50, 1189–1195 (2018).

27. M. Sausen et al., Clinical implications of genomic alterations in the tumour and circulation of pancreatic cancer patients. Nature communications 6, 7686 (2015).

28. J. B. Dawkins et al., Reduced Expression of Histone Methyltransferases KMT2C and KMT2D Correlates with Improved Outcome in Pancreatic Ductal Adenocarcinoma. Cancer research 76, 4861–4871 (2016).

29. C. G. Simone et al., Characteristics and outcomes of adenosquamous carcinoma of the pancreas. Gastrointestinal cancer research: GCR 6, 75–79 (2013).

30. K. Yamaguchi, M. Enjoji, Adenosquamous carcinoma of the pancreas: a clinicopathologic study. Journal of surgical oncology 47, 109–116 (1991).

31. Q. Sun et al., Competition between human cells by entosis. Cell research 24, 1299–1310 (2014).

32. C. de la Cova, M. Abril, P. Bellosta, P. Gallant, L. A. Johnston, Drosophila myc regulates organ size by inducing cell competition. Cell 117, 107–116 (2004).

33. C. Claveria, G. Giovinazzo, R. Sierra, M. Torres, Myc-driven endogenous cell competition in the early mammalian embryo. Nature 500, 39–44 (2013).

34. J. C. Hamann et al., Entosis Is Induced by Glucose Starvation. Cell reports 20, 201–210 (2017).

## References

1. H. L. Mackay et al., Genomic instability in mutant p53 cancer cells upon entotic engulfment. Nature communications 9, 3070 (2018).

2. A. Dobin et al., STAR: ultrafast universal RNA-seq aligner. Bioinformatics (Oxford, England) 29, 15–21 (2013).

3. P. G. Engstrom et al., Systematic evaluation of spliced alignment programs for RNA-seq data. Nature methods 10, 1185–1191 (2013).

4. R. A. Moffitt et al., Virtual microdissection identifies distinct tumor- and stroma-specific subtypes of pancreatic ductal adenocarcinoma. Nature genetics 47, 1168–1178 (2015).

5. A. Subramanian et al., Gene set enrichment analysis: a knowledge-based approach for interpreting genome-wide expression profiles. Proceedings of the National Academy of Sciences of the United States of America 102, 15545–15550 (2005).

6. S. Oki et al., ChIP-Atlas: a data-mining suite powered by full integration of public ChIP-seq data. EMBO Rep, (2018).

7. P. Shannon et al., Cytoscape: a software environment for integrated models of biomolecular interaction networks. Genome research 13, 2498–2504 (2003).

8. A. P. Makohon-Moore et al., Limited heterogeneity of known driver gene mutations among the metastases of individual patients with pancreatic cancer. Nature genetics 49, 358–366 (2017).

9. J. G. Reiter et al., Minimal functional driver gene heterogeneity among untreated metastases. Science 361, 1033–1037 (2018).

10. H. Li, R. Durbin, Fast and accurate short read alignment with Burrows-Wheeler transform. Bioinformatics (Oxford, England) 25, 1754–1760 (2009).

11. M. A. DePristo et al., A framework for variation discovery and genotyping using next-generation DNA sequencing data. Nature genetics 43, 491–498 (2011).

12. L. E. Mose, M. D. Wilkerson, D. N. Hayes, C. M. Perou, J. S. Parker, ABRA: improved coding indel detection via assembly-based realignment. Bioinformatics (Oxford, England) 30, 2813–2815 (2014).

13. K. Cibulskis et al., Sensitive detection of somatic point mutations in impure and heterogeneous cancer samples. Nature biotechnology 31, 213–219 (2013).

14. R. Shen, V. E. Seshan, FACETS: allele-specific copy number and clonal heterogeneity analysis tool for high-throughput DNA sequencing. Nucleic acids research 44, e131 (2016).

15. C. J. Tokheim, N. Papadopoulos, K. W. Kinzler, B. Vogelstein, R. Karchin, Evaluating the evaluation of cancer driver genes. Proceedings of the National Academy of Sciences of the United States of America 113, 14330–14335 (2016).

16. T. Davoli et al., Cumulative haploinsufficiency and triplosensitivity drive aneuploidy patterns and shape the cancer genome. Cell 155, 948–962 (2013).

17. M. S. Lawrence et al., Mutational heterogeneity in cancer and the search for new cancer-associated genes. Nature 499, 214–218 (2013).

18. A. V. Biankin et al., Pancreatic cancer genomes reveal aberrations in axon guidance pathway genes. Nature 491, 399–405 (2012).

19. N. Waddell et al., Whole genomes redefine the mutational landscape of pancreatic cancer. Nature 518, 495–501 (2015).

20. Cancer_Genome_Atlas_Research_Network, Integrated Genomic Characterization of Pancreatic Ductal Adenocarcinoma. Cancer Cell 32, 185–203 e113 (2017).

21. J. T. Robinson et al., Integrative genomics viewer. Nature biotechnology 29, 24–26 (2011).

22. C. M. Bielski et al., Genome doubling shapes the evolution and prognosis of advanced cancers. Nature genetics 50, 1189–1195 (2018).

23. J. G. Reiter et al., Reconstructing metastatic seeding patterns of human cancers. Nature communications 8, 14114 (2017).

